# Nanotherapy-mediated biased antigen-presenting cell reprogramming supports T cell anergy to prevent type 1 diabetes

**DOI:** 10.64898/2026.10.21.760780

**Authors:** Natalie R. Klug, El Hadji Arona Mbaye, Szumo Wang, Sultan Almunif, C. Ramsey Grace, Joseph W. Reda, Débora B. Scariot, Simseok A. Yuk, Yu-Gang Liu, Drew H. Landrowski, Swagat Sharma, Haeik Park, Jeffrey A. Hubbell, Clive H. Wasserfall, Evan A. Scott, Jacqueline A. Burke

**Affiliations:** Department of Biomedical Engineering, Northwestern University, Evanston, IL, USA, 60208; Querrey Simpson Institute for Regenerative Engineering, Northwestern University, Chicago, IL, USA, 60611; Bioengineering Institute, King Abdulaziz City for Science and Technology, Riyadh, Saudi Arabia, 11442; Department of Pathology, Immunology, and Laboratory Medicine, University of Florida, Gainesville, FL, USA, 32610; Departments of Chemical and Biomolecular Engineering and of Biomedical Engineering, New York University, New York, NY, USA, 11021; Department of Biomedical Engineering, NanoSTAR Institute, University of Virginia School of Medicine, Charlottesville, VA, USA, 22904; Departments of Biochemistry and Chemistry, New York University, New York, NY, USA, 10012; Department of Biochemistry and Molecular Pharmacology, NYU Langone Health, New York, NY, USA, 10016

**Keywords:** type 1 diabetes (T1D), disease stage, antigen-agnostic immune tolerance, nanotherapy, T cell anergy, antigen-presenting cells (APCs)

## Abstract

Engineering immune tolerance using endogenously expressed autoantigens could enable early intervention in autoimmune disease. Present therapies suppress T cells to slow disease progression. Here, we develop rapamycin-loaded poly(ethylene glycol)-*b*-poly(propylene sulfide) polymersomes (rPS) to modulate antigen-presenting cells (APCs) during disease-associated antigen presentation. The rPS morphology enables lymphatic drainage following subcutaneous administration to access APCs in draining lymph nodes. Treatment reduced APC costimulatory marker coexpression and shifted CD4⁺ T cells toward an FR4^hi^ CD73^hi^ anergic phenotype. In a type 1 diabetes (T1D) nonobese diabetic (NOD) mouse model, rPS significantly delayed disease onset relative to the standard of care and selectively prevented hyperglycemia. Mediation via APCs, rPS avoids adverse effects including transient lymphopenia. These findings demonstrate how biomaterial-directed drug delivery can reshape APC–T cell interactions, supporting an antigen-agnostic strategy for delaying autoimmune diabetes without systemic lymphocyte depletion.

**The bigger picture:** Type 1 diabetes (T1D) results from a complex autoimmune attack on insulin-producing pancreatic β cells. While the detection of islet autoantibodies before T1D onset enables the development of early intervention strategies, the temporal dynamics of islet antigen presentation and the full repertoire of relevant islet antigens remain unclear. Current immunotherapies primarily target T cells to halt islet injury but can cause transient systemic immune suppression and are not curative. Herein, we identify a potential new therapeutic strategy by which antigen-presenting cells (APCs) are reprogrammed to arrest T1D progression without systemic lymphodepletion. To this end, we utilize rapamycin-loaded polymersome (rPS) nanocarriers which selectively deliver the rapamycin payload to APCs. When given prior to disease onset, rPS treatment delays and prevents hyperglycemia in comparison to a T cell-directed, antibody-mediated systemic therapy.

Importantly, our study provides a comprehensive evaluation of therapeutic intervention and disease progression. Through subcutaneous injection, the nanotherapeutic approach leverages lymphatic drainage to reach the draining lymph nodes. Here, rPS-induced costimulation-deficient APCs can prevent T cell activation without supplying a specific antigen in prediabetic mice. Complex immunoprofiling validated APC reprogramming in vivo and characterized T cell activation state in presymptomatic mice. Overall, this novel biomaterial-mediated approach demonstrates the importance of designing early intervention therapies to disrupt the T cell activation pathway and offer protection to functional β cells and an opportunity to achieve long-term insulin independence.

**Highlights:**

- Rapamycin-loaded polymersomes (rPS) are an APC-directed nanotherapy to slow T1D progression
- A short course of subcutaneous (SC) rPS therapy delays T1D onset in early prediabetic mice
- APC-mediated reprogramming halts T1D progression without lymphodepletion
- Maintenance of effector-memory CD4+ T cell anergy is critical to prevent hyperglycemia

**Graphical Abstract:** 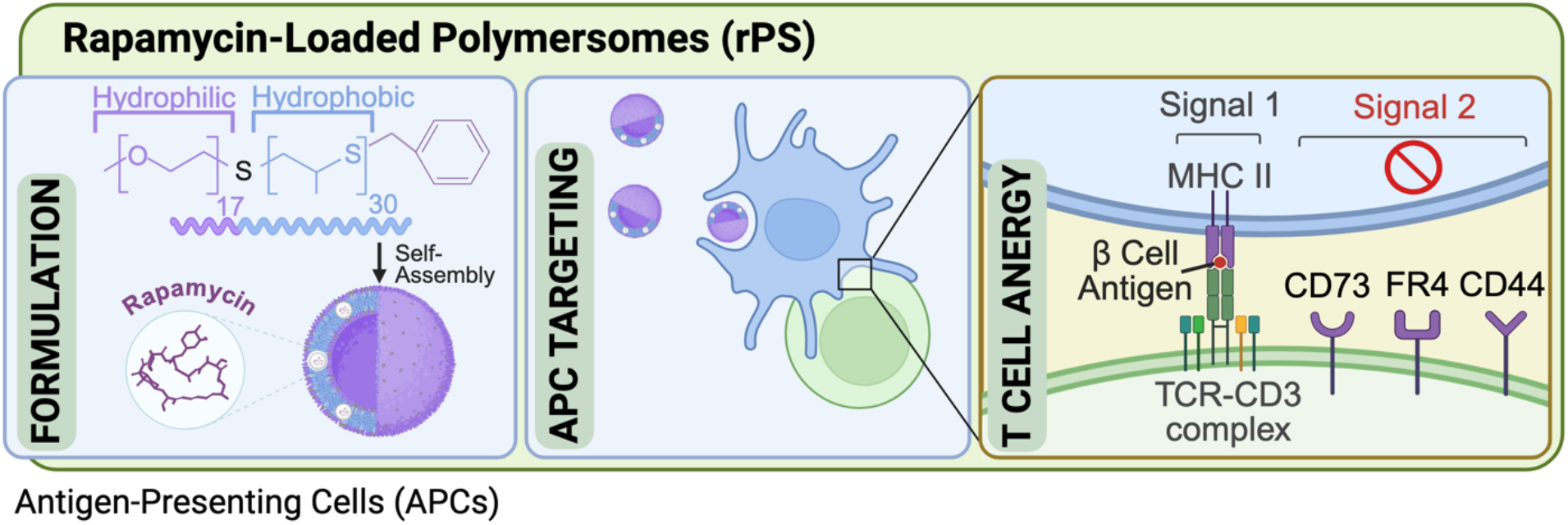

## Introduction

Type 1 diabetes (T1D) is an autoimmune disorder resulting from T cell-mediated destruction of insulin-producing β cells within the pancreatic islets of Langerhans (1). T1D requires lifelong exogenous insulin therapy and carries significant morbidity, including acute complications such as severe hypoglycemia and diabetic ketoacidosis, and chronic complications including nephropathy, neuropathy, retinopathy, and cardiovascular diseases (2). Advances in detecting autoantibodies against β cells enable identification of at-risk individuals before clinical diagnosis, creating an opportunity for preventative immunotherapies to protect β cells (3,4).

Teplizumab (anti-CD3 mAb) is the first and only FDA-approved disease-modifying therapy for delaying β cell damage associated with T1D (5–9). In at-risk individuals, anti-CD3 mAb delayed diabetes onset by 2 to 3 years (8–10). In recently diagnosed patients, anti-CD3 mAb reduced the decline of C-peptide, a biomarker of endogenous insulin production (6). This antigen-agnostic therapy binds to a cell surface antigen receptor present on naïve T cells and interferes with associated signaling, but anti-CD3 mAb therapy is not curative and causes transient lymphodepletion (7). Therefore, next-generation therapies must establish durable islet tolerance while maintaining robust immunity.

Alternatively, pre-clinical approaches aim to induce antigen-specific tolerance by either delivery of disease-relevant antigens or ex vivo expansion and transfusion of regulatory T cells (Tregs) (11–16). However, substantial interpatient heterogeneity in the autoantigens contributing to disease progression may limit efficacy of islet-specific antigen delivery (17). Additionally, Treg expansion requires individualized cell isolation and customized expansion, making broad clinical implementation infeasible (18). We previously showed that rapamycin-loaded polymersomes (rPS), formed from poly(ethylene glycol)_17_-b-poly(propylene sulfide)_30_ (PEG_17_-b-PPS_30_) copolymer, altered the organ and cellular level biodistribution of rapamycin by facilitating delivery to lymph node resident APCs following subcutaneous (SC) injection (19–21). In a fully major histocompatibility (MHC) mismatched mouse model of islet transplantation, rPS induced costimulation-deficient APCs, leading to CD4+ T cell depletion and anergy as well as CD8+ Treg expansion within the draining lymph nodes (LNs) (19). At 100 days post-transplant, splenic T cells isolated from SC rPS-treated recipient mice demonstrated significantly reduced proliferation in response to stimulation with allogeneic donor antigen; however, T cells maintained their ability to respond to a foreign allogeneic antigen (19). These results may also suggest that rPS therapy could induce antigen-specific immunomodulation in vivo, without ex vivo cell modification or antigen delivery.

In this work, we hypothesized that reprogramming T cells in vivo via modulation of antigen presenting cell (APC) signaling in the presence of native antigen within secondary lymphoid organs could induce T1D delay. After SC injection, PEG-b-PPS nanocarriers are primarily taken up by monocytes in draining LNs, enabling access to systemic antigen repertoires (19,22,23). Given the ability of rPS to induce antigen-specific islet tolerance for allogeneic transplant, we evaluated the ability of this nanotherapy to preserve functional native β cell mass by restoring immune tolerance to islet autoantigens in the spontaneous nonobese diabetic (NOD) mouse model. Despite limitations in translating findings from this model to the clinical setting (24), it is the principal preclinical model for evaluating autoimmune responses contributing to the development of T1D onset (25–27). In this model, T1D develops spontaneously in a sex and colony-specific manner. The median onset age is 18 to 22 weeks for female NOD mice (28). To study the rapamycin dose-response, while controlling for nanocarrier number, we developed rPS formulations with constant polymer content and increasing rapamycin payloads. We benchmarked the efficacy of SC rPS formulations in delaying diabetes onset against intravenous (IV) administration of anti-CD3 mAb to prediabetic NOD mice. Next, we used spectral flow cytometry to assess mechanistic differences between APC- and T cell-targeted immunomodulation by rPS and anti-CD3 mAb, respectively. The optimized SC rPS regimen induced tolerogenic APCs and maintained effector-memory CD4+ T cells in an anergic state to preserve islet tolerance. Overall, our findings identify APC-directed nanotherapy as a disease-modifying strategy for arresting T1D onset through restoration of immune tolerance without broad immunosuppression. Antigen agnostic APC targeted therapy has the potential to be applied broadly to autoimmune diseases.

## Methods

### Mice

Female, 7-week-old NOD/ShiLtJ mice were procured from the Jackson Laboratory and housed in a pathogen-free facility within the Center for Comparative Medicine at Northwestern University (NU). All experiments were performed in accordance with NU IACUC-approved protocols.

### Chemicals

Unless specifically stated, all chemicals were from Sigma Aldrich.

### Polymer Synthesis

Nanocarriers were fabricated from PEG_17_-b-PPS_30_ block copolymer as previously described (19). PEG_750_-OH was modified with a mesylate group, then functionalized to an intermediate PEG_750_-thioacetate so the thiolate anion could induce ring opening polymerization of propylene sulfide. Benzyl bromide end-capping yielded PEG_17_-b-PPS_30_-Bz. Polymer chain length and formation of the diblock copolymer were confirmed via ^1^H-NMR and gel permeation chromatography.

### Nanocarrier Formulation and Characterization

Formulations were made via thin film rehydration as previously described (19,29). PEG_17_-b-PPS_30_-Bz (20 mg) was dissolved with dichloromethane. For rPS formulations, rapamycin (0.25, 0.5, or 1 mg; Selleckchem; dissolved at 25 mg/mL in pure ethanol) was added. The solution was vortexed and then desiccated overnight to remove organic solvents. The resulting polymer film was rehydrated with sterile DPBS (1 mL) and shaken (1500 rpm, 24 hours) to allow particle formation.

Unencapsulated drug was removed using Sephadex LH20 columns in sterile DPBS. Particle size distribution was measured by dynamic light scattering (DLS). Nanoparticle tracking analysis (NTA) via the NanoSight300 was used to confirm particle size and quantify particles concentrations. Small angle X-ray scattering (SAXS) measurements were performed on the 5-ID-D beamline at the Dow-Northwestern-DuPont Collaborative Access Team (DND-CAT) of the Advanced Photon Source, Argonne National Laboratory. Data collection settings are in the Supplementary. The spherical, vesicular nanocarrier morphology was verified via cryogenic transmission electron microscopy (cryo-TEM). Samples were vitrified using a Leica EM GP2 plunge freezer with backside blotting. Lacey copper grids (300 mesh) were glow-discharged for 45 seconds prior to sample application. After blotting from the grid’s reverse side for 4 seconds using filter paper, grids were plunged into liquid ethane. Cryo-TEM data was collected on a Thermo Fisher Glacios transmission electron microscope operating at 200 kV, equipped with a Falcon 4 direct electron detector. Images were acquired at a nominal magnification of 120 kX, corresponding to a pixel size of 1.2 Å/pixel.

Previously published high-performance liquid chromatography methods were used to quantify rapamycin concentration in rPS formulations (19). Endotoxin was quantified via chromogenic Limulus Amebocyte Lysate assay (GenScript).

### Assessment of Diabetes Incidence in NOD Mice

Prediabetic 8 or 12-week-old female NOD mice served as untreated controls or received rPS SC (6 doses, Q3D) or mouse anti-CD3 mAb (clone 145-2C11, BioXCell) IV (5 doses, QD, 5 µg/dose). All SC doses were administered by scruff injections, and all IV doses were administered via tail-vein infusions. For the 8-week intervention study, 3 doses of rPS were evaluated: rPS Low_Rapa_ (0.0625 mg/mL rapamycin, 0.5 mg rapamycin/kg/dose), rPS Medium_Rapa_ (0.125 mg/mL rapamycin, 1 mg rapamycin/kg/dose), and rPS High_Rapa_ (0.250 mg/mL rapamycin, 2 mg rapamycin/kg/dose). For the 12-week intervention study, rPS Medium_Rapa_ (0.125 mg/mL rapamycin, 1 mg rapamycin/kg/dose), in addition to controls vehicle PS SC and rapamycin PO (0.125 mg/mL rapamycin, 1 mg rapamycin/kg/dose; VistaPharm) was assessed. The polymer concentration in each formulation was constant at 6.7 mg/mL. Blood glucose (Supplementary Figs. 1 and 2) and weight were monitored TIW. T1D onset was determined by two consecutive readings >250 mg/dL recorded on separate days. Upon onset, mice were removed from the study. Whole blood was collected retro-orbitally for CBC analysis. Serum was separated for insulin autoantibody (IAA) detection. One-month post-treatment, mice were fasted for 12 hours before an oral glucose tolerance test. A bolus injection of a 30% glucose solution (2 g/kg) was administered, and blood glucose was recorded at designated time points for 120 minutes (Supplementary Fig. 3).

### IAA Detection

IAA was measured using an Islet Autoantibody Standardization Program (IASP)-certified luciferase immunoprecipitation assay adapted by the Wasserfall Lab (30,31). Briefly, serum was incubated overnight at 4°C with luciferase-tagged proinsulin. Immunocomplexes were captured by Protein A&G Sepharose beads (Cytiva). Nano-Glo luciferase substrate (Promega) was added, and relative light units (RLU) were measured on a luminometer (Berthold). Samples exceeding a preliminary cut-off (RLU ≥ 8074) were run in a competition assay: serum was incubated with luciferase-labeled proinsulin and 10.5 ng unlabeled insulin (Humulin R, Lilly). Delta RLUs were calculated (ΔRLU = RLUnon-compete – RLUcompete) to determine the IAA positivity cut-off (ΔRLU ≥ 5049).

### Histology

Harvested mouse pancreas were fixed in 10% neutral buffered for 4 hours and stored in in 70% ethanol. Subsequently, samples were paraffin-embedded, sectioned (6 sections per pancreas, 4 µm thick, 150 µm apart) and stained with hematoxylin & eosin (H&E) and insulin (C27C9 rabbit mAb, Cell Signaling Technology). A NanoZoomer 2.0-HT series captured whole slide images (20X). Islet and pancreatic area were measured using NDP2.view. Insulitis scoring was completed using H&E staining in accordance with standard guidelines (Supplementary Fig. 4) (25,32).

### CBC Analysis

Whole blood samples were run on Element HT5 analyzer yielding a 5-part differential, including values for white blood cells (WBC; Supplemental Fig. 5).

### Flow Cytometry

Whole blood and A/B LN were collected, processed into single cell suspensions and stained with fluorochrome-conjugated antibodies (see Supplementary Table 1 and 2), according to previously published methods (19,21,29). Samples were acquired on an Aurora spectral flow cytometer (Cytek) with spectral unmixing via SpectroFlo (Cytek), then analyzed using FlowJo 2. The gating strategy is outlined in Supplementary (Supplementary Figs. 6-8, Supplementary Tables 3-12).

## Statistics

Data represented as mean ± standard deviation (SD). All statistical analyses were completed in GraphPad Prism10. Mantel-Cox log-rank survival statistics were employed for median T1D onset. One- and two-way ANOVAs with Tukey’s multiple comparison test were used to compare differences between rPS and other treatment groups unless stated otherwise.

## Data and Resources Availability

All datasets generated and needed to evaluate and reproduce results in this paper are present in the paper and/or the Supplementary Materials.

## Results

### Generation and characterization of rPS with varied concentrations of rapamycin

We hypothesized that selective delivery of rapamycin to APCs via rPS would lead to controlled immunomodulation in vivo. With a constant payload, the number of nanoparticles can impact delivery and therefore efficacy of the therapeutic (33,34). To properly assess a range of rapamycin doses, we prepared rPS formulations with constant polymer content, particle number and physiochemical properties (size, charge) while varying the rapamycin amount loaded into the PS hydrophobic membrane (Fig. 1*A*). In our previous work, a 6-dose regimen of rPS (0.125 mg/mL rapamycin, 1 mg rapamycin/kg/dose) prevented islet graft rejection (19). Therefore, we compared this reference rPS formulation with formulations encapsulating two-fold higher (rPS High_Rapa_, 0.250 mg/mL rapamycin) or two-fold lower (rPS Low_Rapa_, 0.0625 mg/mL rapamycin) rapamycin concentrations. Increasing the rapamycin payload did not alter particle size or vesicular morphology (Fig. 1*B* and *C*). Additionally, all rPS formulations exhibited similar drug loading efficiencies, particle size, particle concentration, and zeta potentials. All formulations contained less than 0.05 endotoxin units (EU)/mL, the endotoxin threshold established by the United States Pharmacopeia for preclinical studies (Fig. 1*D*) (35). Cryo-TEM) images confirmed the bilayer membrane structure and vesicle morphology of each rPS formulation (Fig. 1*E*-*G*). Thus, increasing the amount of rapamycin in each polymersome nanocarrier did not alter particle integrity.

**Fig. 1.**
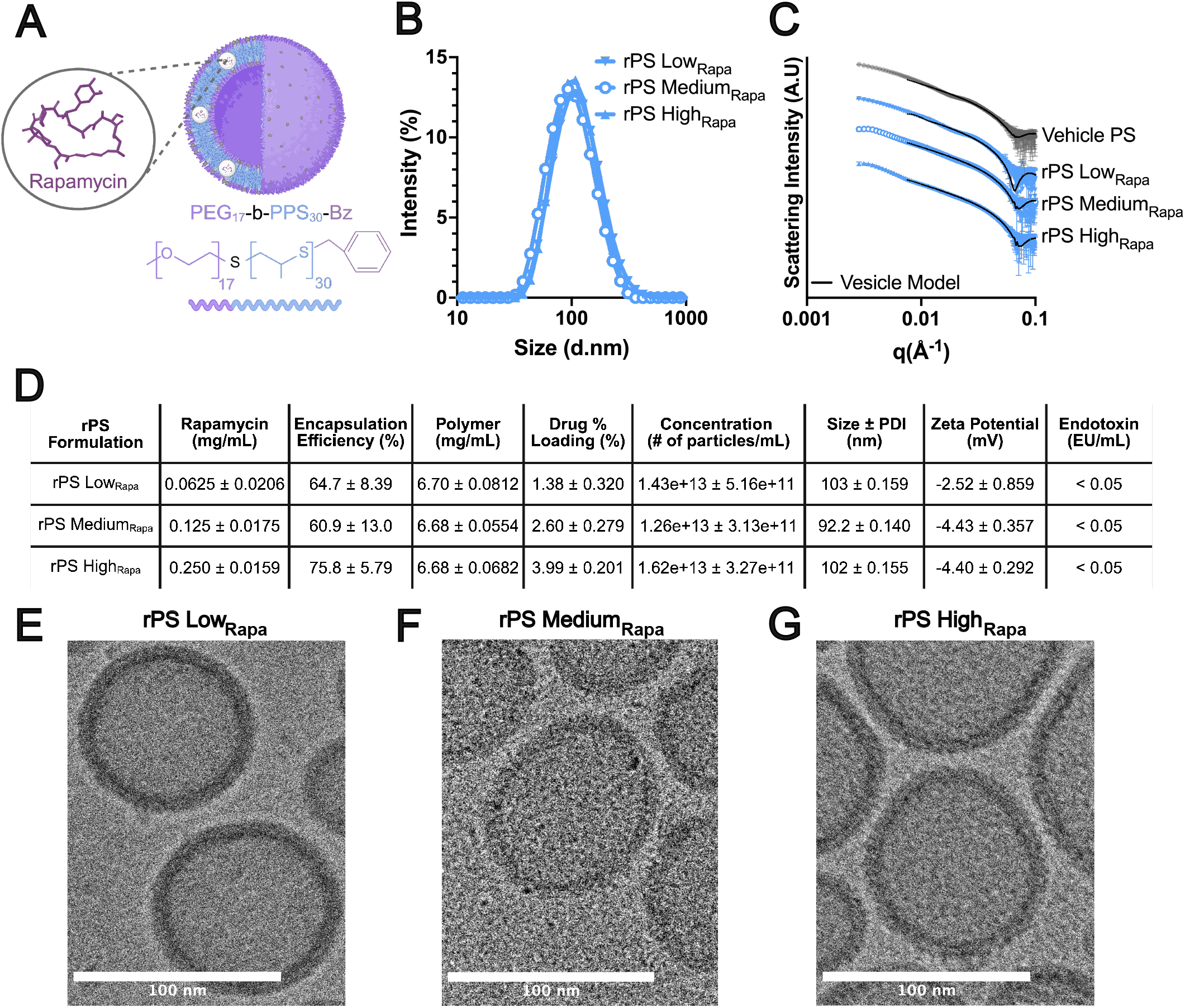
Engineering polymersomes with graded rapamycin payloads. *A*: Schematic of rapamycin loading in the hydrophobic membrane of PEG_17_-b-PPS_30_-Bz polymersomes. rPS formulations contained consistent polymer content and variable rapamycin payload. *B*: Size distribution of rPS formulations as measured by DLS. *C*: Modeling of SAXS scattering profiles confirmed the vesicular morphology of rPS formulations. *D*: Summary of rPS characterization. Data presented as mean ± SD. *E*-*G*: Cryo-TEM images confirm the vesicular particle morphology with a hydrophobic membrane for all rPS formulations (scale bar = 100 nm). These formulations supported the subsequent investigations.

### SC rPS delays T1D onset in prediabetic NOD mice

From our previous work in islet transplantation (19), we hypothesized that administering rPS to prediabetic NOD mice would reprogram APCs towards a tolerogenic phenotype and therefore regulate T cells to protect native β cell mass. However, the dynamics of antigen presentation in allogeneic islet transplantation and autoimmune diabetes onset differ (26,36). Therefore, we assessed various dosing schedules, doses, and intervention timepoints to better understand the pharmacodynamics of rPS in prediabetic NOD mice. Based on single and 3-dose pilot studies in prediabetic NOD mice (Supplementary Fig. 1) and the effective 6-dose regimen used in the allogeneic islet transplantation mouse model (19), we selected an rPS dosing regimen consisting of 6 doses, given every third day. We intervened at 8 weeks of age, before the expansion of effector T cells but after initial T cell priming had occurred (26). We routinely monitored mice for hyperglycemia. Although anti-CD3 mAb was only effective only in reversing newly established diabetes in NOD mice, we compared SC rPS therapy with the IV dosing regimen previously evaluated for delaying T1D onset in prediabetic NOD mice (5 μg/day for 5 consecutive days) (*n* = 16) (37,38). Here, we explored 3 different rapamycin doses of rPS, using the formulations described in Fig. 1. Equivalent volumes of rPS Low_Rapa_ (0.0625 mg rapamycin/mL), rPS Medium_Rapa_ (0.125 mg rapamycin/mL) and rPS High_Rapa_ (0.250 mg rapamycin/mL) delivered doses of 0.5, 1.0, and 2.0 mg rapamycin/kg/dose, respectively (Fig. 2*A*). The study classified T1D onset as early or late according to whether a mouse developed T1D before or after 21 weeks of age (Fig. 2*A*) (39). In this study, the median onset age for non-treated mice was 21.7 weeks (Fig. 2*B*). Anti-CD3 mAb slowed early T1D onset but the mice had a similar median onset age of 21.4 weeks (Fig. 2*B*). All rPS doses significantly delayed onset in comparison to anti-CD3 mAb mice (*p = 0.0078, 0.0063, and 0.0175*, respectively). The rPS High_Rapa_-treated mice reached a median onset age of 28.6 weeks and were the only rPS-treated group to reach 50% onset before 30 weeks of age (Fig. 2*B*). In the rPS Medium_Rapa_ and rPS Low_Rapa_ treated groups, less than 50% of mice developed diabetes by the 30-week study endpoint. rPS Medium_Rapa_ delayed onset 10 weeks longer than the non-treated (*p = 0.0493)* and anti-CD3 mAb-treated (*p = 0.0063)* groups. Similarly, rPS Low_Rapa_ delayed onset 10 weeks longer than the non-treated (*p = 0.129)* and anti-CD3 mAb-treated (*p = 0.0078)* groups. Furthermore, half of the non-treated mice developed early onset T1D (50%, 8/16 mice) and under a quarter (18.8%, 3/16 mice) developed late onset T1D. With anti-CD3 mAb treatment, 43.8% (7/16 mice) of mice developed early onset T1D and 37.5% (6/16 mice) showed late onset. Overall, rPS resulted in the lowest proportion of mice that developed T1D before 21 weeks; no mice in the rPS High_Rapa_ group and only one mouse each in the rPS Medium_Rapa_ and rPS Low_Rapa_ groups developed early T1D (6.25%, 1/16 mice) (Fig. 2*D* and Supplementary Fig. 1). While the proportion of rPS-treated mice that developed late onset T1D was equal to that of the anti-CD3 mAb-treated mice (37.5%, 6/16 mice), most rPS Medium_Rapa_-treated mice remained diabetes-free at 30 weeks of age (56.3%, 9/16 mice).

**Fig. 2.**
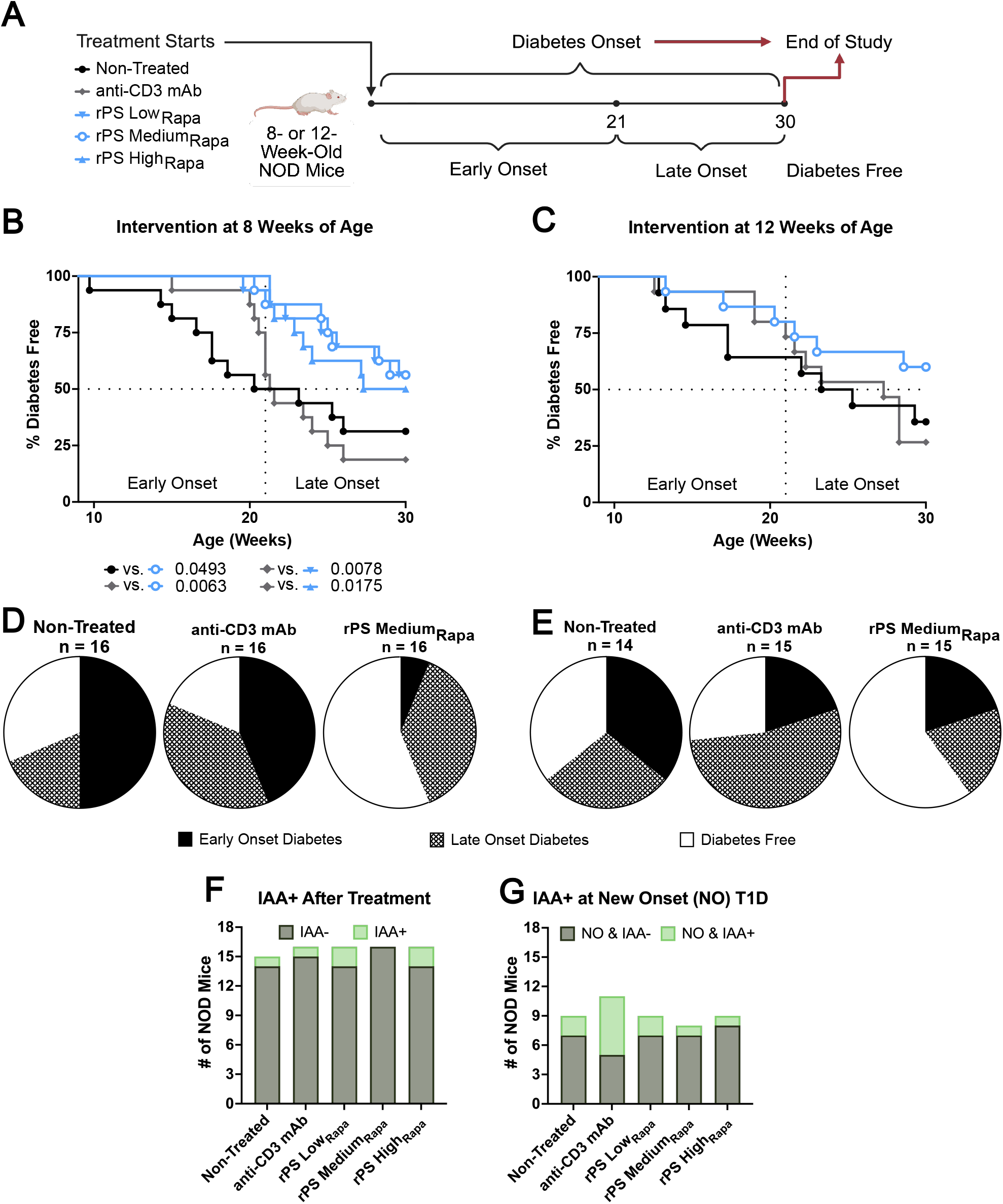
rPS delays T1D onset in prediabetic NOD mice. *A*: Schematic of experimental design. *B* and *C*: Diabetes incidence in female NOD mice following intervention at (*B*) 8 weeks of age with IV anti-CD3 mAb (5 μg, QD, 5 doses) or SC rPS (Low_Rapa_ at 0.5 mg/kg, Medium_Rapa_ at 1 mg/kg, or High_Rapa_ at 2 mg/kg; Q3D, 6 doses) or (*C*) 12 weeks of age with IV anti-CD3 mAb (5 μg, QD, 5 doses) or SC rPS (Medium_Rapa_ at 1 mg/kg, Q3D, 6 doses) benchmarked against non-treated mice (*n* = 14-16). The vehicle-control group was omitted from the 8-week intervention study based on prior validation (19) to reduce animal use and prioritize the primary treatment comparisons. Log-rank tests generated the P-values shown for the 8-week intervention cohort in the Kaplan-Meier curve. *D* and *E*: Proportion of mice that developed early onset diabetes, late onset diabetes or remain diabetes-free at 30 weeks of age following intervention at 8 (*D*) or12 (*E*) weeks of age. *F* and *G*: Serum IAAs measured 24 hours post-treatment and at new onset (NO) (8-week intervention cohort). No association with onset by chi-squared with Bonferroni correction.

To assess the effect of intervention timing, mice received the rPS Medium_Rapa_ dosing regimen beginning at 12 weeks of age, closer to disease onset (*n* = 14-15) (Fig. 2*C* and Supplementary Fig. 2). Early preclinical studies found that anti-CD3 mAb more effectively delayed diabetes onset in older NOD mice (12 weeks or 14 to 25 weeks of age), with presumably severe insulitis (37,38). With intervention starting at 12 weeks of age, anti-CD3 mAb-treated mice achieved median onset at 27.3 weeks, slowing the time to onset by 3 weeks compared to non-treated mice with median onset at 24.3 weeks (Fig. 2*C*). As with intervention at 8 weeks of age, more than 50% of rPS Medium_Rapa_-treated mice remained diabetes-free through 30 weeks of age, precluding calculation of median disease onset. Over 50% of anti-CD3 mAb-treated mice developed late onset T1D (53.3%, 8/15). While there was no significant difference between the median onset ages for the treatment groups in this study, only rPS Medium_Rapa_-treated mice had an increase in the number of diabetes-free mice at 30 weeks (60%, 9/15). In summary, across both the 8- and 12-week cohorts, most non-treated mice developed early onset T1D, whereas more anti-CD3 mAb-treated mice developed late onset T1D, and the majority of rPS Medium_Rapa_-treated mice remained diabetes-free (Fig. 2*D* and *E* and Supplementary Figs. 1-4).

The detection of autoantibodies against β cells, including IAAs, is associated with islet infiltration in both humans and NOD mice (27,40). We collected serum from the 8-week cohort upon completion of treatment and upon confirmation of sustained hyperglycemia. The luciferase immunoprecipitation (LIPS) assay quantified serum IAAs (30). IAA detection did not increase immediately following any treatment regimen, and IAA positivity remained low across all groups (Fig. 2*F*). T1D onset, IAA detection, and therapeutic intervention showed no significant correlations (Fig. 2*G*) (27).

### SC rPS avoids transient lymphodepletion in NOD mice

Clinically, anti-CD3 mAb infusions cause transient lymphodepletion during dosing (5,7). Additionally, anti-CD3 mAb can induce mild cytokine release syndrome (CRS) during treatment, leading to symptoms such as weight loss due to increasing metabolic demands (41). Building on our previous findings, we hypothesized that SC rPS would enable more precise control over immunomodulation in the draining LNs via APCs, rather than causing broad immunosuppression in the blood (19). To test the systemic effects of rPS, we monitored peripheral blood counts and body weight change during treatment (Fig. 3*A*). Anti-CD3 mAb-treated mice experienced a significant loss in weight midway through treatment (*p < 0.0001* versus non-treated, *p = 0.0075* versus rPS Low_Rapa_, *p = 0.0026* versus Medium_Rapa_) (Fig. 3*B*). Complete blood count (CBC) analysis showed that 37.5% of the anti-CD3 mAb-treated mice developed leukopenia (Fig. 3*C*) and 100% developed lymphopenia (Fig. 3*D*). Similarly, anti-CD3 mAb led to a transient decrease in neutrophils and monocytes in the blood (*p = 0.0004* and *p = 0.0009* in comparison to non-treated) (Supplementary Fig. 5). Notably, the severe white blood cell depletions for the anti-CD3 treated group occurred midway through dosing, coinciding with the significant decrease in weight (Fig. 3*B*). Conversely, rPS did not reduce leukocytes, lymphocytes, neutrophils, monocytes or body weight during treatment.

**Fig. 3.**
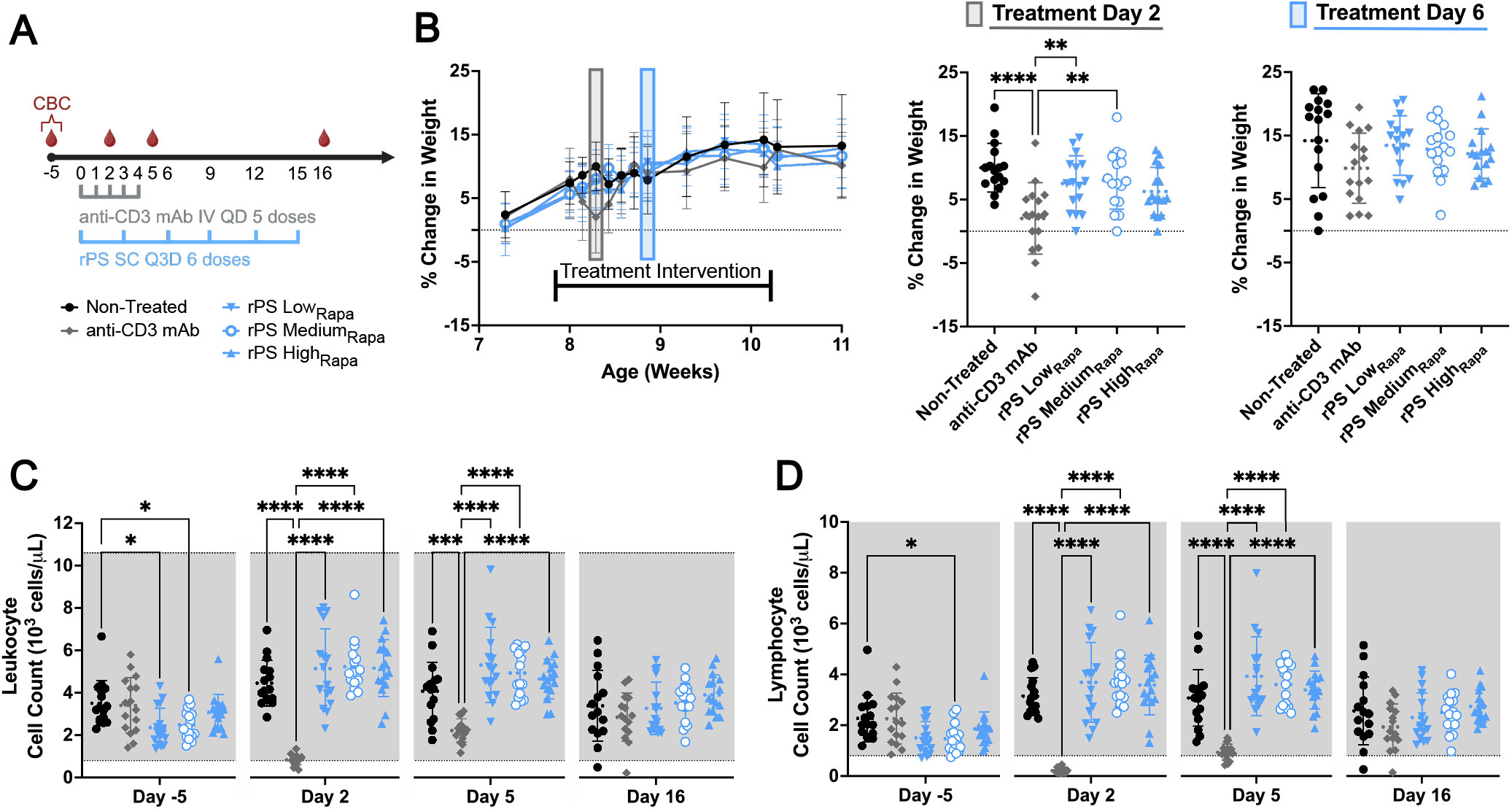
rPS treatment does not cause transient lymphodepletion. *A*: Schematic of experimental design for CBC sample collections at baseline, halfway through each treatment, and 24-hours after the final treatment for each group. *B*: Average percent change in weight, calculated from baseline, during treatment (*n* = 16). Bracket defines treatment intervention window. Grey and blue boxes designate the mid-treatment point for anti-CD3 mAb (Day 2, middle panel) and rPS treatments (Day 6, right panel). Statistical significance determined by one-way ANOVA followed by Tukey’s multiple-comparisons test. \*\**P* < 0.01, \*\*\*\**P* < 0.0001. *C*: Leukocyte count over time and by treatment. *D*: Lymphocyte count over time and by treatment. Grey shaded area represents the reference range. All statistical analyses performed using mixed-effects model followed by Tukey’s multiple-comparisons test. \**P* < 0.05, \*\*\**P* < 0.001, \*\*\*\**P* < 0.0001. All data represented as mean ± SD.

### SC rPS conditions APC-mediated CD4+ T cell anergy in vivo

To probe the mechanism of action that enables rPS-induced tolerance, 8-week-old NOD mice (*n* = 4-5) received the 6-dose rPS Medium_Rapa_ regimen (Figs. 2*A*, 3*A*), and immunophenotyping characterized specific cellular populations 24 hours after the completion of all treatment regimens. Here, mice received oral rapamycin, SC vehicle PS, IV anti-CD3 mAb, SC rPS Medium_Rapa_, or remained non-treated. Instead of a systemic administration route like oral and IV, the SC PS platform leverages lymphatic drainage to access APCs and facilitate trafficking to the lymph nodes closest to the site of injection (Supplementary Fig. 8) (4,19,20). Therefore, the nontoxic PS platform facilitates APC uptake and reprogramming dependent on the loaded cargo. Similar to prior work, initial in vitro studies confirmed that rapamycin encapsulation altered costimulatory marker expression on APCs (Supplementary Fig. 9). Now, we used spectral flow cytometry to assess the costimulatory and coinhibitory marker expression on the APCs in the draining LNs (axial/brachial, A/B) and blood (Supplementary Fig. 10) immediately after treatment completion. A heatmap displays the relative frequencies of APC subsets, normalized to the average non-treated control (Fig. 4*A*). rPS lowered the frequency of CD40+, CD86+, PD-L1+, MHC I+, and MHC II+ APCs, including B cells, dendritic cells (DCs), and monocyte-and-macrophage lineage cells (M/Ms) compared to anti-CD3 mAb-treated mice. Following rPS treatment, there was a decrease in the frequency of DCs (of CD45+ cells) (Fig. 4*B*), and specifically a reduction of activated and mature CD40+ CD86+ PD-L1+ MHC I+ MHC II+ DCs (Fig. 4*C* and *D* and Supplementary Fig. 11). The frequency of M/Ms increased after anti-CD3 mAb treatment (Fig. 4*E*) while rPS again reduced costimulatory CD40+ CD86+ PD-L1+ MHC I+ MHC II+ M/Ms (Fig. 4*F* and *G* and Supplementary Fig. 11). Anti-CD3 mAb also showed an increased B cell frequency after treatment (Supplementary Fig. 12). Furthermore, rPS elicited costimulation-deficient APCs, shifting the LN cellular composition towards a tolerogenic environment.

**Fig. 4.**
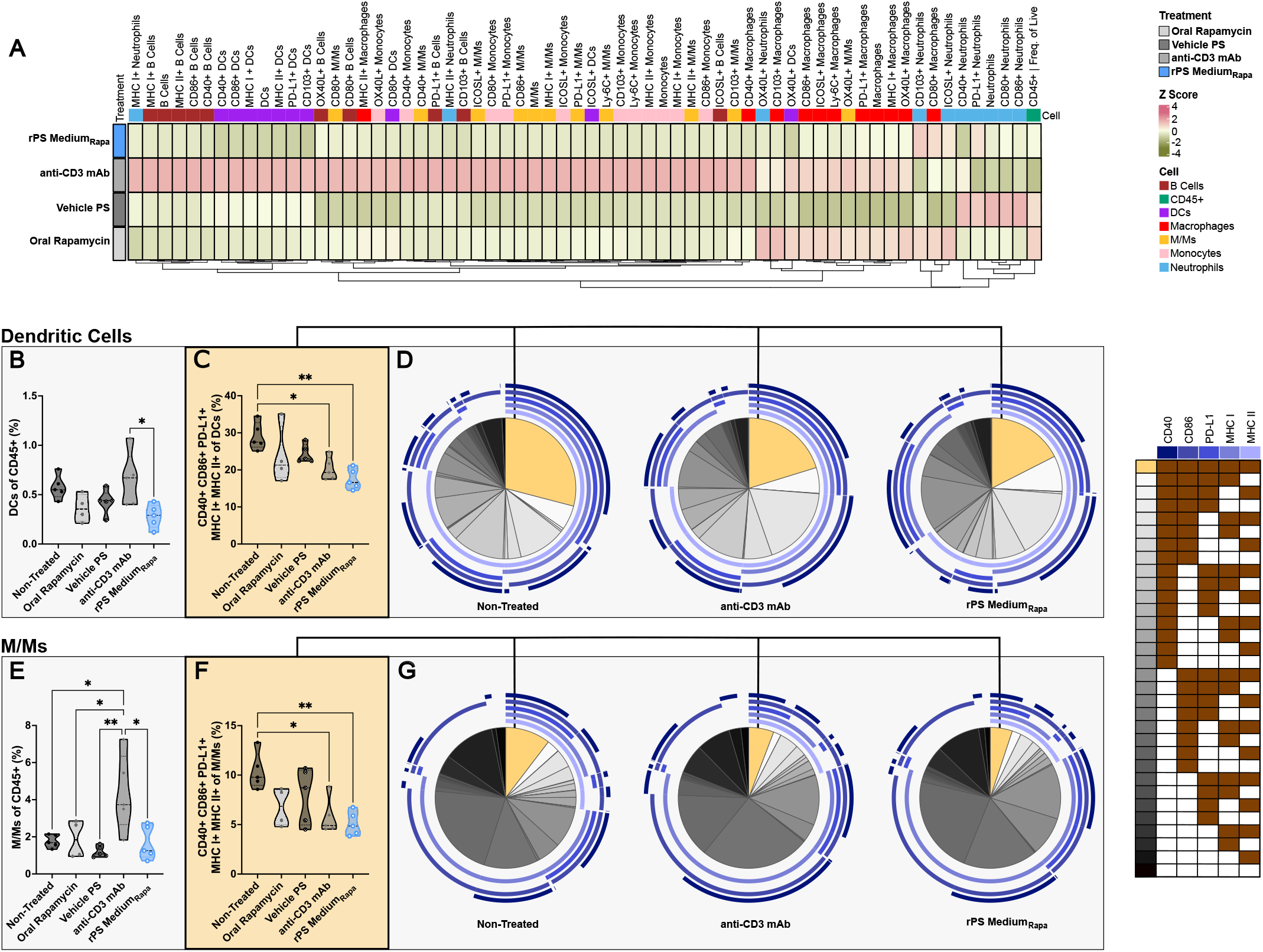
rPS treatment induces costimulatory-deficient APCs in vivo. Flow cytometry analysis performed on the A/B LN collected from non-treated NOD mice or NOD mice treated with oral rapamycin (1 mg rapamycin/kg, Q3D, 6 doses), SC polymersomes (Q3D, 6 doses), IV anti-CD3 mAb (5 μg, QD, 5 doses), or SC rPS Medium_Rapa_ (1 mg rapamycin/kg, Q3D, 6 doses), starting at 8 weeks of age (*n* = 4-5). Study completed on A/B LNs, harvested 24 hours after treatment ended. *A*: The analysis normalized flow cytometry data (CD45+) to the non-treated group, calculated Z scores, clustered samples by Euclidean distance, and visualized the results as a heatmap, where each column represents a treatment group and each row represents frequency of a specific phenotype. *B* and *E*: Percentage of dendritic cells (DCs) (*B*) or monocyte- and-macrophage lineage cells (M/Ms) (*E*) of CD45+ cells. (*C* and *F*) Percentage of CD40+ CD86+ PD-L1+ MHC I+ MHC II+ DCs (*C*) or M/Ms (*F*). All data are presented as mean ± SD with \**P* < 0.05, \*\**P* < 0.01. Statistical significance determined by one-way ANOVA followed by Tukey’s multiple-comparison test. *D* and *G*: Pie charts show distribution of mean receptor co-expression by DCs (*D*) and M/Ms (*G*) for non-treated, anti-CD3 mAb, and rPS Medium_Rapa_ groups. Yellow slice corresponds to combination shown in *C* and *F*.

In parallel, spectral flow cytometry quantified activation and anergy marker expression on CD4+ T cells isolated from the A/B LNs (Supplementary Fig. 13). As in humans, MHC II-associated genetic susceptibility predisposes NOD mice to develop T1D by promoting the activation of CD4+ T cells, which help coordinate CD8+ T cell cytotoxic effector function (42–44). We therefore hypothesized that the rPS-induced costimulation-deficient APC phenotype would alter the CD4+ T cell compartment, thereby disrupting the proliferation and differentiation of both CD4+ and CD8+ T cells. The gating strategy outlined in Fig. 5*A*-*C* defined CD4+ T cells subsets (44–46). Naïve CD4+ T cells (CD44-FoxP3-) were separated from CD4+ regulatory T cells (Tregs, CD44± FoxP3+) (Fig. 5*A*). FR4 and CD73 coexpression further resolved effector-memory CD4+ T cells (CD44+ FoxP3-) into activated effector-memory (T_EM_, FR4^lo^ CD73^lo^) and anergic (T_anergic_, FR4^hi^ CD73^hi^) phenotypes (Fig. 5*B* and *C*) (44,45,47). From these definitions, anti-CD3 mAb caused a significant decrease in naïve CD4+ cells (Fig. 5*D*). In response, the frequency of CD4+ Tregs increased following anti-CD3 mAb treatment. This result aligns with the proposed mechanism of tolerance induced by anti-CD3 mAb where the reduction of naïve T cells is rebalanced by the regulatory population (5,48). Alternatively, rPS decreased activated T_EM_ cells and induced the hyporesponsive T_anergic_ phenotype (FR4^hi^ CD73^hi^) of T_EM_ cells (Fig. 5*E* and *F*). A heatmap compares the relative T cell frequencies in response to treatment, normalized to the control non-treated group (Fig. 5*G*). The heatmap illustrates that rPS and anti-CD3 mAb modulated distinct T cell populations in the A/B LN. A tSNE visualization reveals these populations of CD4+ T cells in the A/B LN (Fig. 5*H* and *I*). Anti-CD3 mAb expanded activated and suppressive FR4+, ICOS+, and ICOS+ CD38+ CD4+ T_EM_ populations (Fig. 5*J* and *K*). In addition, anti-CD3 mAb led to an expansion of highly suppressive ICOS+ FR4+ CD4+ Tregs (Fig. 5*L* and *M*). While mice treated with anti-CD3 mAb experienced an overall reduction in naïve CD4+ T cells, activated FR4+ and CD38+ CD4+ T cells predominantly remained (Fig. 5*N*-*P*). Thus, anti-CD3 mAb influenced a broad shift of T cell subsets towards suppression while rPS maintained T cells in an anergic, tolerogenic state.

**Fig. 5.**
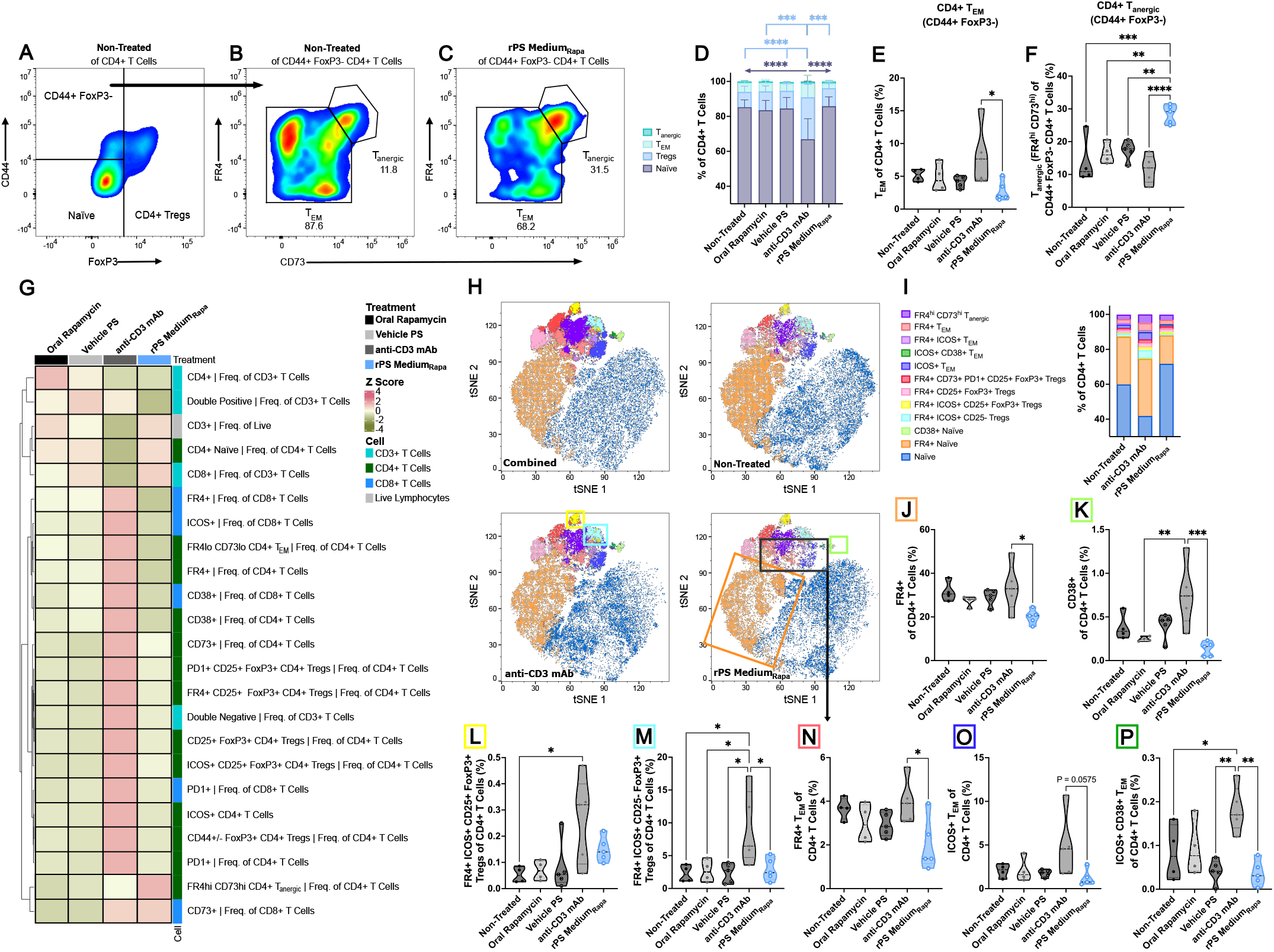
rPS-mediated costimulation deficient APCs maintain T cell anergy. The study performed flow cytometry analysis on the A/B LNs collected from non-treated NOD mice or NOD mice treated with oral rapamycin (1 mg rapamycin/kg, Q3D, 6 doses), SC polymersomes (Q3D, 6 doses), IV anti-CD3 mAb (5 μg, QD, 5 doses), or SC rPS Medium_Rapa_ (1 mg rapamycin/kg, Q3D, 6 doses), starting at 8 weeks of age (*n* = 4-5). All A/B LNs harvested 24 hours after treatment ended. *A*-*C*: Gating strategy for naïve, CD4+ regulatory T cells (Tregs), effector-memory T cells (T_EM_), and anergic T cells (T_anergic_). *D*: Distribution of CD4+ T cell subpopulations by treatment group. \*\*\**P* < 0.001, \*\*\*\**P* < 0.0001 using two-way ANOVA followed by Tukey’s multiple-comparisons test. ***E***: Percentage of T_EM_ of CD4+ T cells. *F*: Percentage of T_anergic_ of CD44+ FoxP3-CD4+ T cells. \**P* < 0.05, \*\**P* < 0.01, \*\*\**P* < 0.001, \*\*\*\**P* < 0.0001 using one-way ANOVAs followed by Tukey’s multiple-comparisons test. *G*: The analysis normalized the flow cytometry data for T cells (CD3+) to the non-treated group, calculated Z scores, clustered samples by Euclidean distance, and visualized the results as a heatmap, where each column represents a treatment group and each row represents frequency of a specific phenotype. *H*: tSNE visualization of CD4+ T cells by treatment. *I*: Distribution of CD4+ T cell phenotypes identified in the tSNE plots. *J* and *K*: Percentage of FR4+ (*J*) and CD38+ (*K*) naïve T cells. *L* and *M*: Percentage of FR4+ ICOS+ CD25+ (*L*) and FR4+ ICOS+ CD25-(*M*) Tregs. (*N*-*P*), Percentage of FR4+ (*N*), ICOS+ (*O*), ICOS+ CD38+ (*P*) T_EM_ cells. \**P* < 0.05, \*\**P* < 0.01, \*\*\**P* < 0.001 using one-way ANOVAs followed by Tukey’s multiple-comparisons test.

## Discussion

Our findings support a mechanism in which selective modulation of APCs reprogram APC-T cell interactions to maintain autoreactive T_EM_ cells in an anergic state without inducing lymphodepletion. In contrast to anti-CD3 mAb, which relies on transient T cell depletion and immunosuppression (41,48), rPS supports immune tolerance by reducing APC costimulatory signaling and promoting hyporesponsive T cell fates without inducing T cell exhaustion. Together, our data identify APC-directed induction of T cell anergy as a distinct mechanism for preventing immune diabetes. By exploiting endogenous antigen presentation rather than requiring disease-specific antigen delivery or individualized Treg therapies, this work identifies a fundamentally independent strategy for antigen-agnostic tolerance with broad translational potential across autoimmune diseases. Our data demonstrate that rPS establishes both prolonged delay and selective prevention of autoimmune diabetes. Unlike other antigen-specific therapies that require prior knowledge of the target autoantigen (12,15,16,49), APC-directed nanotherapy leverages endogenous antigen presentation to induce tolerance without restricting the immune response to a predefined antigen repertoire. This feature may be particularly advantageous as autoimmune responses evolve through epitope spreading (25), enabling newly revealed islet antigens to be presented in a tolerogenic rather than immunogenic context. Consistent with this concept, diabetes-free survival declined rapidly in anti-CD3 mAb-treated mice after approximately 21 weeks of age, where most rPS-treated mice remained protected throughout the study (25). These findings suggest that reprogramming APC function may provide an enhanced strategy for maintaining immune tolerance as autoimmune disease evolves.

Early therapeutic intervention maximizes the preservation of functional β cell mass and improves long-term disease outcomes (4,8,50). Consistent with previous reports, anti-CD3 mAb treatment at 12 weeks was associated with late T1D onset, but provided little benefit when mice received treatment at 8 weeks (37), a stage preceding the extensive expansion of pathogenic effector and memory T cells (24–26). Because anti-CD3 mAb primarily modulates activated T cells, its efficacy likely depends on the presence of an established autoreactive T cell population (48). In contrast, rPS maintained efficacy regardless of the timing of intervention, consistent with a distinct mechanism of action. By targeting APCs, rPS acts upstream of effector T cell expansion to attenuate costimulatory signaling and reshape T cell priming before pathogenic immune responses become fully established. This ability to intervene early in disease progression may preserve a larger functional β cell reserve, thereby potentially extending onset delay or enabling prevention.

Notably, this work supports a pathway by which SC rPS-induced tolerogenic APCs regulate the fate of both naïve and T_EM_ cells without causing widespread immunosuppression. In the draining LNs, SC rPS led to a reduction in mature and costimulatory DCs and M/Ms coexpressing CD86, CD40, PD-L1, MHC I, and MHC II. The absence of costimulation limited naïve CD4+ T cell activation or depletion. In addition, rPS-induced APCs sustained FR4^hi^ CD73^hi^ T_anergic_ cells, which are unresponsive to self-antigens due to incomplete costimulation (44,45). In the NOD mouse model, this T_anergic_ phenotype can decline with age and as mice develop diabetes (45). Therefore, rPS can sustain peripheral tolerance through the induction of anergic T_EM_ cells to evade autoimmunity. Importantly, activated DCs and M/Ms advance T1D progression by driving local inflammatory responses, presenting antigens, and further activating pro-inflammatory B cells, which can also lead to islet autoantibody production (50,51). Thus, anti-CD3 mAb allowed for the persistence of activated FR4^lo^ CD73^lo^ T_EM_ cells with enhanced memory phenotypes (FR4+, ICOS+, and ICOS+ CD38+). Furthermore, the residual naïve CD4+ T cells also expressed activated effector T cell markers FR4+ and CD38+ following anti-CD3 mAb treatment. Together, these antigen-experienced cells retain the capacity to differentiate upon antigen re-exposure and could restart the autoimmune response in the presence of costimulatory APCs (26,45). Collectively, these findings establish rPS as a disease-modifying therapy that restores immune tolerance by selectively reprogramming APCs rather than broadly suppressing T cells. By combining efficacy with a clinically practical SC route of administration and without lymphodepletion, this work provides a foundation for translating APC-directed nanotherapy as an early intervention strategy for individuals at risk for developing autoimmune diabetes. To advance to the clinic, future efforts should focus on chemistry, manufacturing, and controls (CMC) considerations for scaling rPS formulations and toxicity studies in more clinically relevant non-human primate models.

## Acknowledgments

The authors would like to thank María Méndez-Santos and Sydney Wiggins for assistance with animal handling and tissue processing, and Hiroaki Sai for the collection and analysis of SAXS data. This work made use of the Keck-II facility (RRID: SCR_026360) of Northwestern University’s NUANCE Center, which has received support from the IIN and Northwestern University’s MRSEC program (NSF DMR-2308691). X-ray scattering studies were performed at the Dow-Northwestern-DuPont Collaborative Access Team (DND-CAT) located at Sector 5 of the Advanced Photon Source (APS) at Argonne National Lab. DND-CAT is supported by Northwestern University, The Dow Chemical Company, and DuPont de Nemours, Inc. This research used resources of the Advanced Photon Source, a U.S. DOE Office of Science User Facility operated for the DOE Office of Science by the Argonne National Laboratory under contract no. DE-AC02-06CH11357. Data was collected using an instrument funded by the National Science Foundation under Award Number 0960140. NTA was performed in the Analytical bioNanoTechnology Equipment Core Facility of the Center for Regenerative Nanomedicine at Northwestern University (RRID:SCR_023706). ANTEC receives support from Feinberg School of Medicine, Northwestern University. Transmission electron micrographs were recorded at the University of Virginia Molecular Electron Microscopy Core facility (RRID: SCR_019031), which is supported in part by the School of Medicine. This work was supported by the Developmental Therapeutics Core at Northwestern University and the Robert H. Lurie Comprehensive Cancer Center support grant (NCI CA060553) (RRID:SCR_017948). Comparative histopathology and molecular phenotyping services were provided by MHPL (RRID:SCR_017870) of Northwestern University which is supported by National Cancer Institute (NCI) P30-CA060553 grant awarded to the Robert H. Lurie Comprehensive Cancer Center. The authors acknowledge the Center for Comparative Medicine Pre-Clinical Services for assistance with clinical pathology sample processing and analysis. We thank the University of Chicago Cytometry and Antibody Technology Facility, especially David LeClerc, for assistance with spectral flow cytometry (RRID: SCR_017760). Figure schematics were created in Biorender.com.

## Author Contributions

N.R.K., E.A.S., and J.A.B. conceived the experiments. N.R.K., E.H.A.M., S.W., S.A., C.R.G., D.B.S., S.A.Y., Y.L., D.H.L., S.S., and H.P. performed the experiments. N.R.K., S.A., C.R.G., and J.W.R. analyzed and visualized the data. J.A.H., C.H.W., E.A.S., and J.A.B. contributed to discussion and/or reviewed the manuscript. N.R.K., E.A.S., and J.A.B. wrote the original draft. N.R.K., J.A.H., E.A.S., and J.A.B. edited the manuscript. J.A.B. is the guarantor of this work and, as such, had full access to all the data in the study and takes responsibility for the integrity of the data and the accuracy of the data analysis.

