## Supplementary Material for "Nanotherapy-mediated biased antigen-presenting cell reprogramming supports T cell anergy to prevent type 1 diabetes"

Natalie Klug *et al.*

**This PDF file includes:**

Supplementary Fig. 1 to 13
Supplementary Data Collection Settings and Methods
Supplementary Tables 1 to 12
Supplementary References

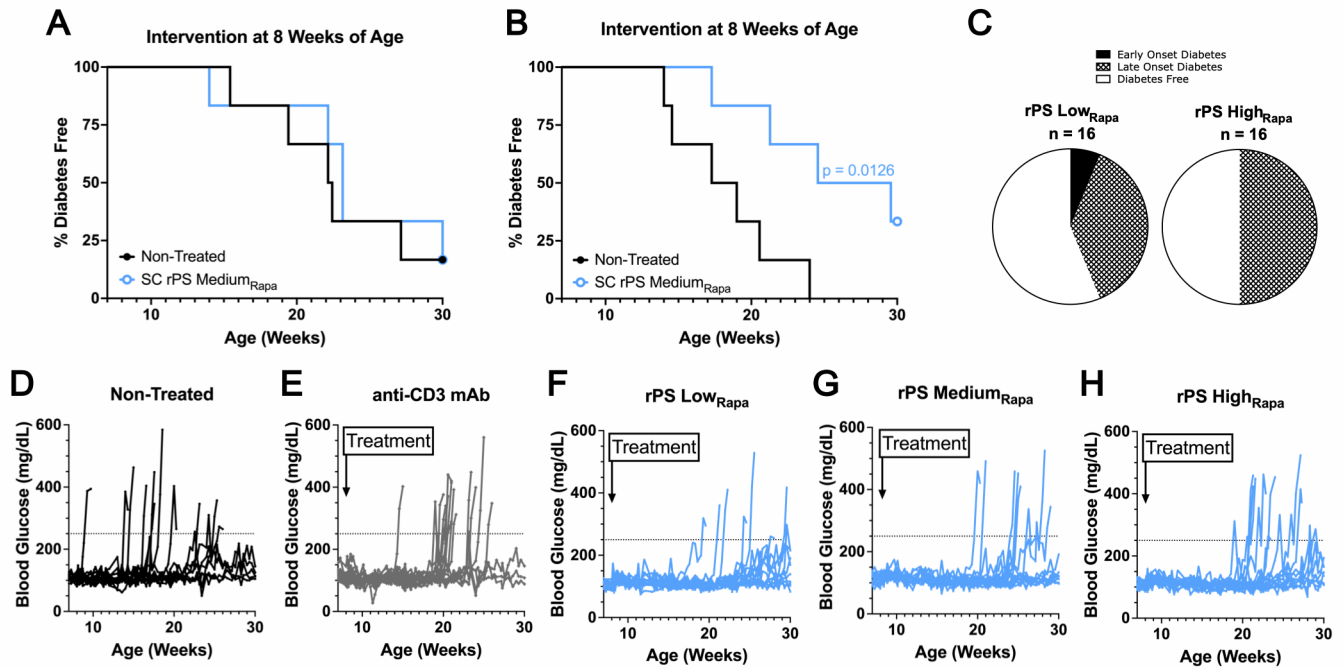

### Supplementary Fig. 1

Development and evaluation of rPS for T1D onset delay in early prediabetic NOD mice. T1D onset determined by two consecutive blood glucose readings above 250 mg/dL, measured on back-to-back monitoring days. *A*: Single dose of SC rPS Medium<sub>Rapa</sub> (1 mg rapamycin/kg) administered to 8-week-old mice ($n = 6$). *B*: Three doses of rPS Medium<sub>Rapa</sub> (1 mg rapamycin/kg) administered to 8-week-old mice every three days ($n = 6$). One non-treated mouse died not due to T1D onset. *C*: Proportion of mice that developed early onset diabetes, late onset diabetes or remain diabetes free at 30 weeks of age following intervention at 8 weeks. *D-H*: Blood glucose trajectories for each mouse dosed at 8 weeks ($n = 16$) shown in Fig. 2B. *I*: Insulin autoantibodies (IAAs) detected at baseline for mice dosed at 8 weeks ($n = 16$) and shown in Fig. 2F-G.

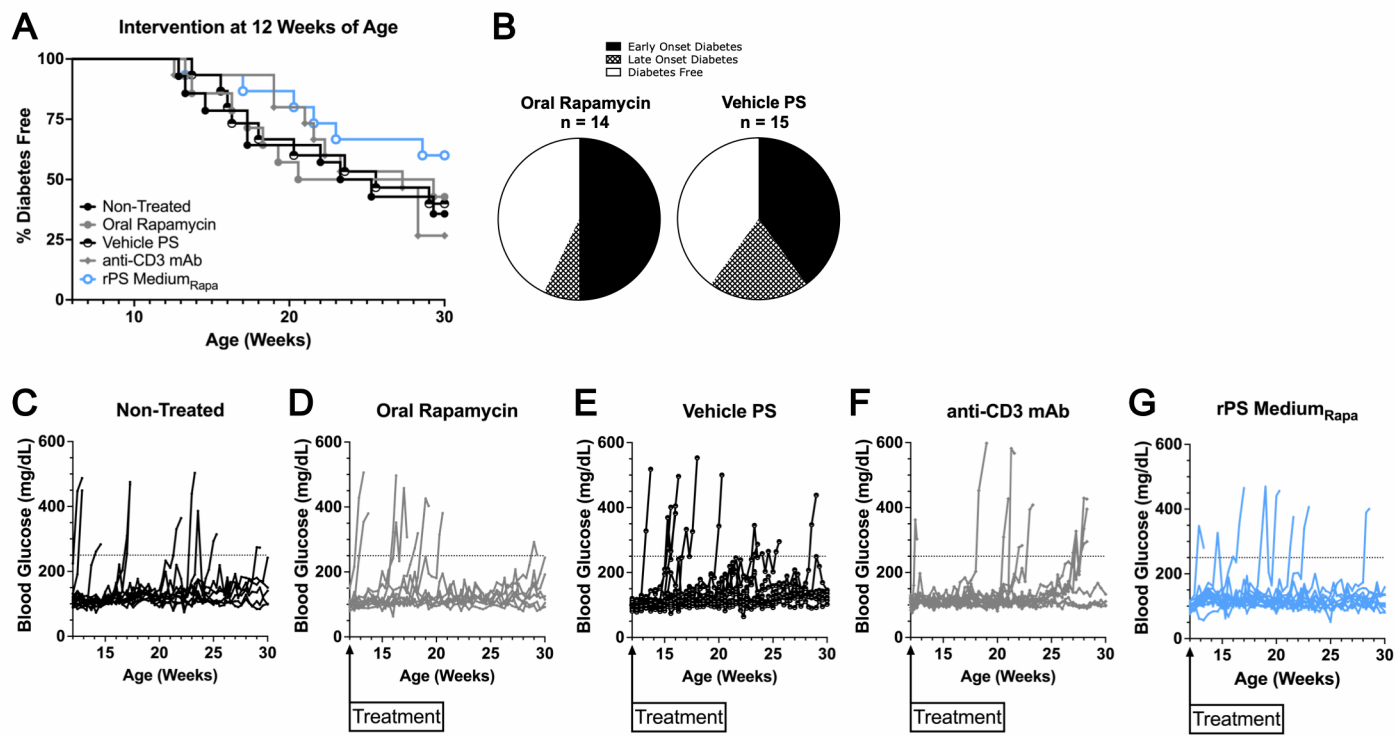

### Supplementary Fig. 2

Development and evaluation of rPS for T1D onset delay in late prediabetic NOD mice. *A*: Six doses of rPS Medium<sub>Rapa</sub> (1 mg/kg of 0.125 mg/mL rPS) administered to 12-week-old mice ($n = 14-15$). rPS Medium<sub>Rapa</sub> treatment benchmarked against six doses of oral rapamycin (1 mg/kg of 0.125mg/mL rapamycin), SC vehicle PS, and IV anti-CD3 mAb (5 μg, 5 days). *B*: Proportion of mice that developed early onset diabetes, late onset diabetes or remain diabetes free at 30 weeks of age following intervention at 12 weeks. *C-G*: Blood glucose trajectories for each mouse dosed at 12 weeks ($n = 14-15$).

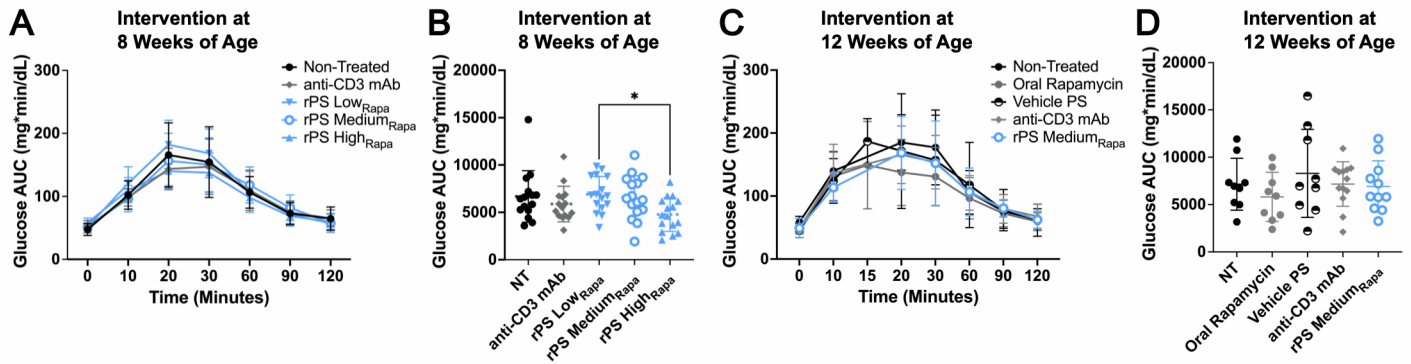

#### Supplementary Fig. 3

Glucose tolerance test in NOD mice treated with rPS. *A* and *B*: Average glucose readings and glucose area under the curve (AUC) during an oral glucose tolerance test (OGTT) administered one month after treatment to normoglycemic mice dosed at 8 weeks ($n = 16$ all groups except $n = 15$ for non-treated). *C* and *D*: Average glucose readings and glucose AUC during an OGTT administered 1 month after treatment to normoglycemic mice dosed at 12 weeks ($n$ varies by group: non-treated $n = 9$, oral rapamycin $n = 9$, vehicle PS $n = 10$, anti-CD3 mAb $n = 14$, and rPS Medium<sub>Rapa</sub> $n = 13$). Statistical analyses completed using one-way ANOVAs followed by Tukey's multiple-comparisons test. $*P < 0.05$.

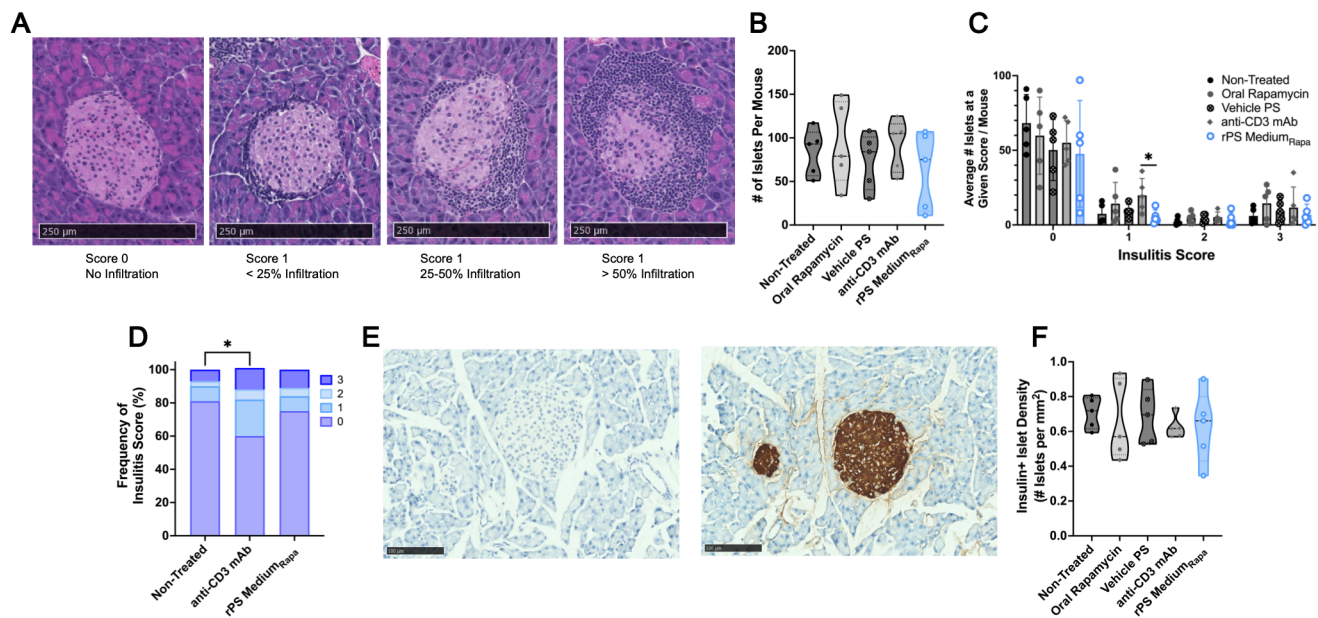

##### Supplementary Fig. 4

Histological analysis of NOD mice treated with rPS. *A*: Insulitis scoring guide. *B* and *C*: Hematoxylin & eosin (H&E) staining analyzed for (*B*) total islet number and (*C*) number of islets at a given insulitis score (right). No significant difference in total number of islets. Statistical analyses completed using Welch's two-tailed t-test on the number of islets with the score of 1. \*$P < 0.05$. *D*: Frequency of each islet score by treatment group. Normalized to total number of islets scored for the respective treatment group. \*$P < 0.05$. Statistical analysis completed using chi-squared analysis with Bonferroni correction for multiple comparison. *E*: Insulin negative and positive staining controls. *F*: Insulin-positive density per acinar area calculated for each treatment group. Statistical analyses completed using one-way ANOVAs followed by Tukey's multiple-comparisons test (not significant).

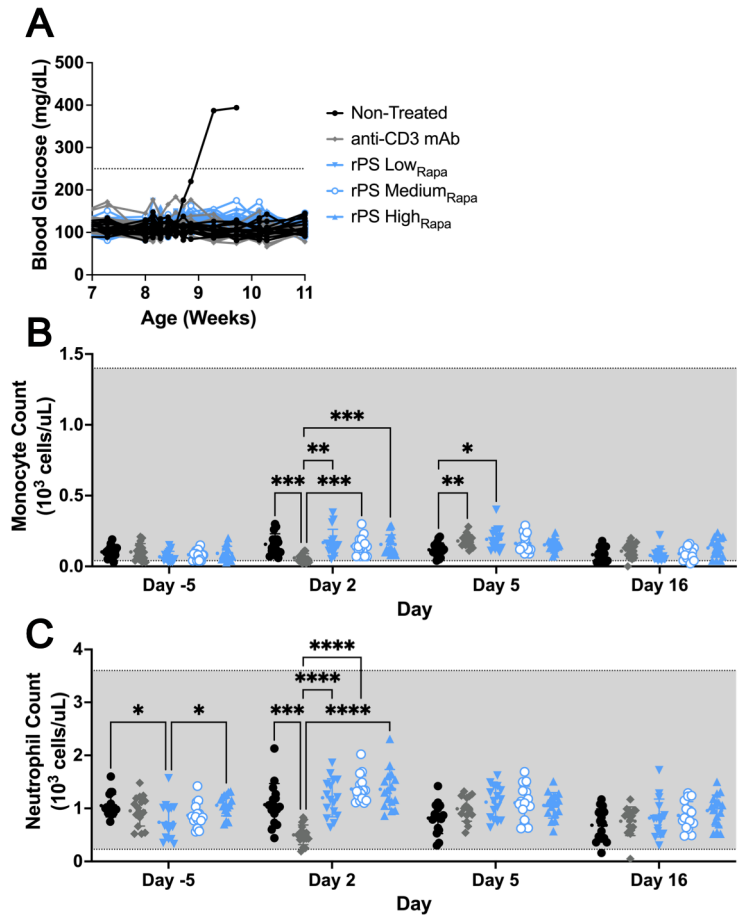

#### Supplementary Fig. 5

rPS SC treatment avoids lymphodepletion during treatment. *A*: Blood glucose recordings during treatment for mice dosed at 8 weeks of age ($n = 16$) from Fig. 2B and Fig. 3. *B*: Change in monocytes over time during treatment. Grey shaded area represents the reference range. *C*: Change in neutrophils over time and by treatment group during treatment. Grey shaded area represents the reference range. All statistical analyses were performed using mixed-effects model followed by Tukey's multiple-comparisons test. $*P < 0.05$, $**P < 0.01$, $***P < 0.001$, $****P < 0.0001$.

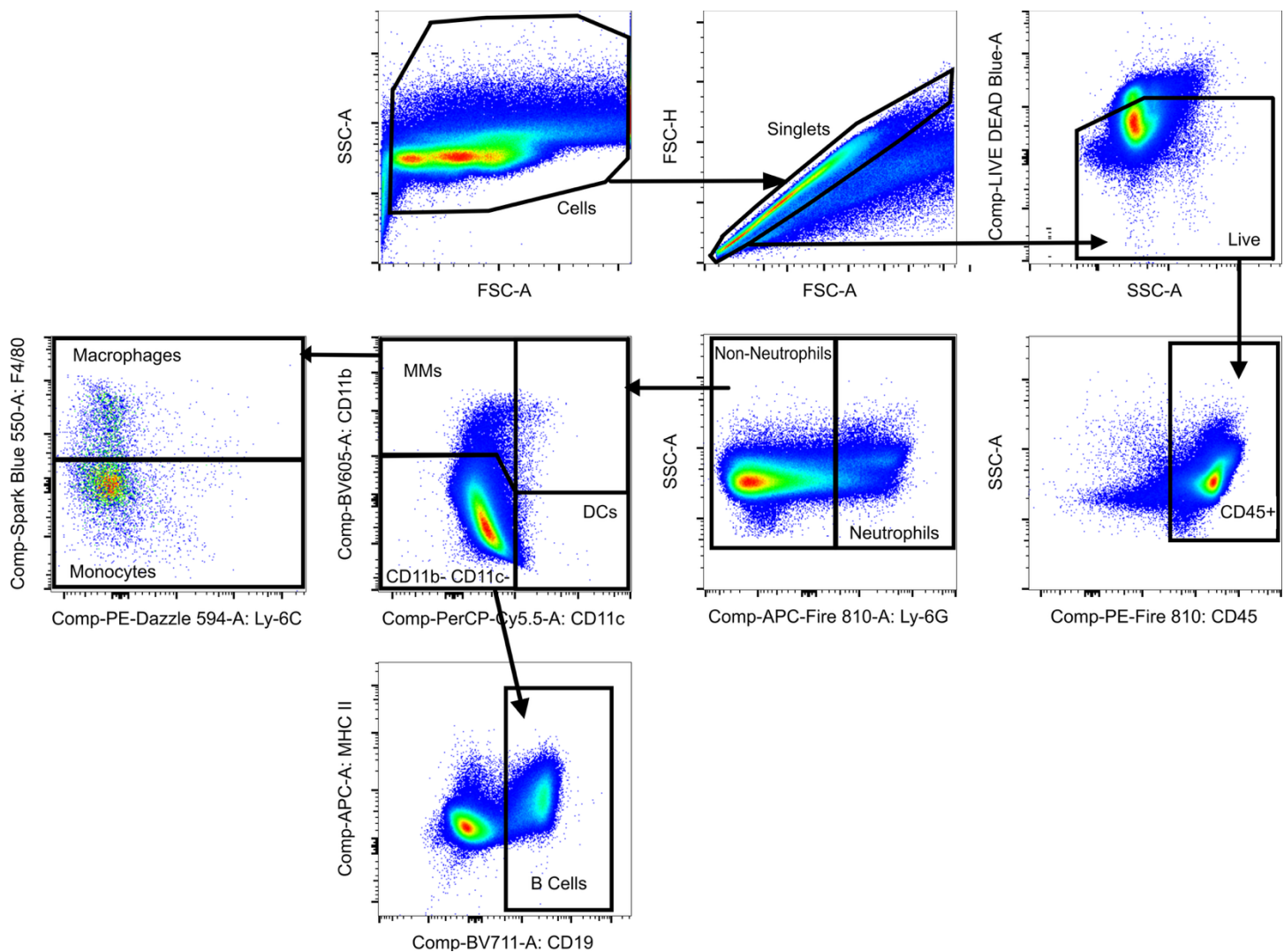

**Supplementary Fig. 6**

Gating strategy for the APC panel. Demonstrated on an A/B LN from a non-treated mouse.

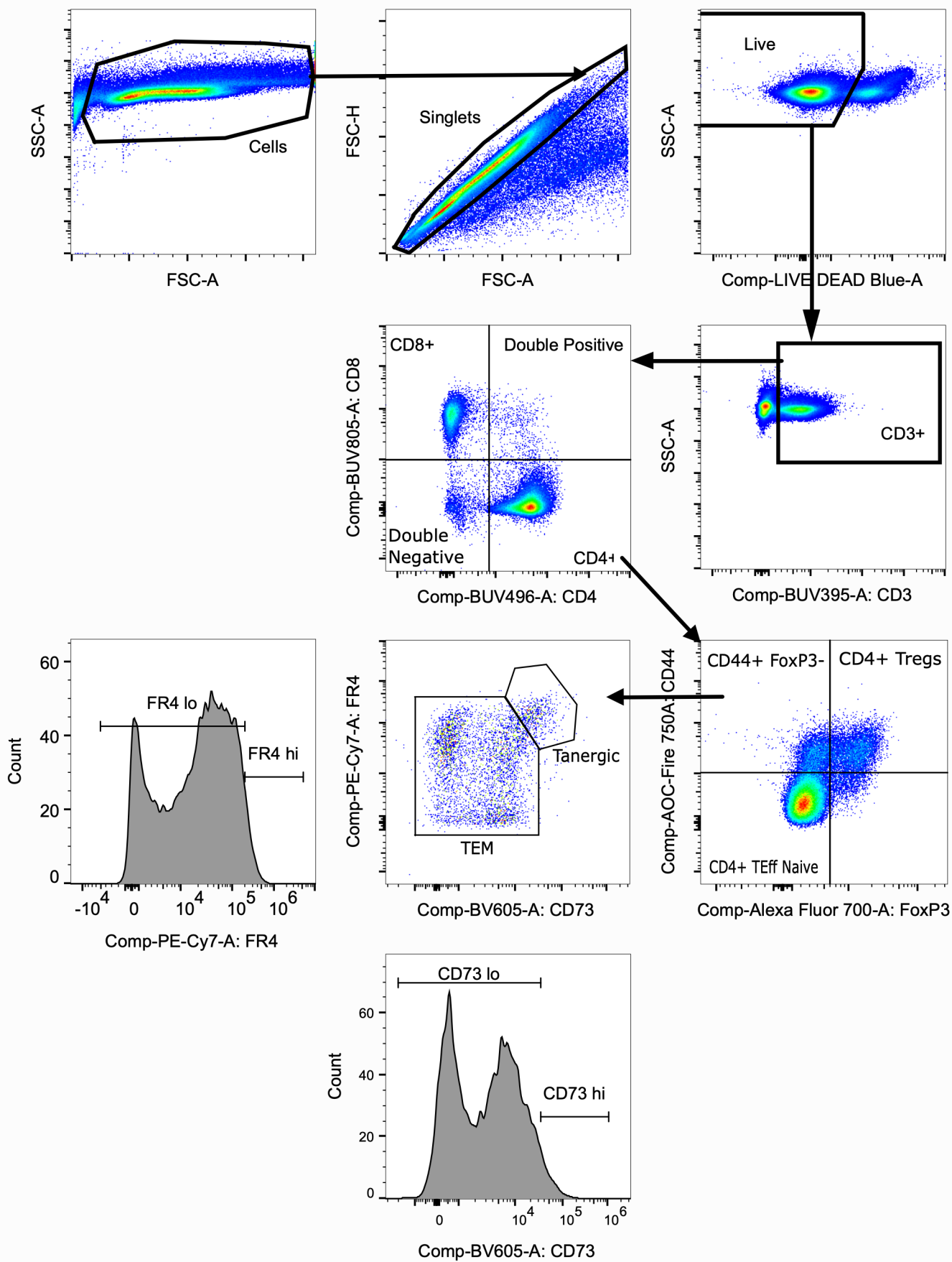

**Supplementary Fig. 7**

Gating strategy for the T cell panel. Demonstrated on an A/B LN from a non-treated mouse.

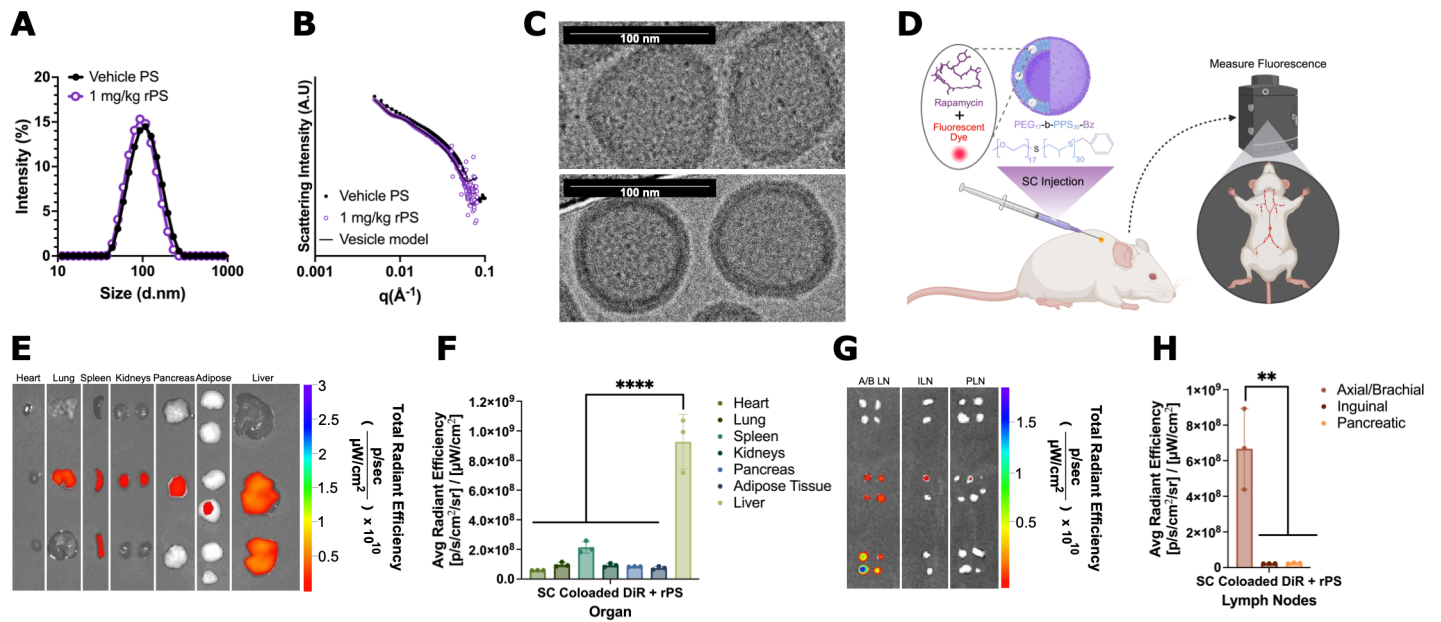

### Supplementary Fig. 8

Generation and biodistribution of polymersome (PS) platform. (A) Dynamic light scattering (DLS) confirmation of PS size. (B) PS morphology aligned with vesicular model fit via small-angle x-ray scattering (SAXS). (C) Representative cryogenic transmission electron microscopy (cryo-TEM) images of PS vesicular morphology (scale bar = 100 nm). (D) SC administration of rapamycin and DiR co-loaded PS. (E-F) Total (E) and average radiant efficiency (F) measured via IVIS for major organs. (G-H) Total (G) and average radiant efficiency (H) measured via IVIS for draining lymph node (LN) populations. All statistical analyses completed using One-Way ANOVA followed by Tukey's multiple-comparisons test.

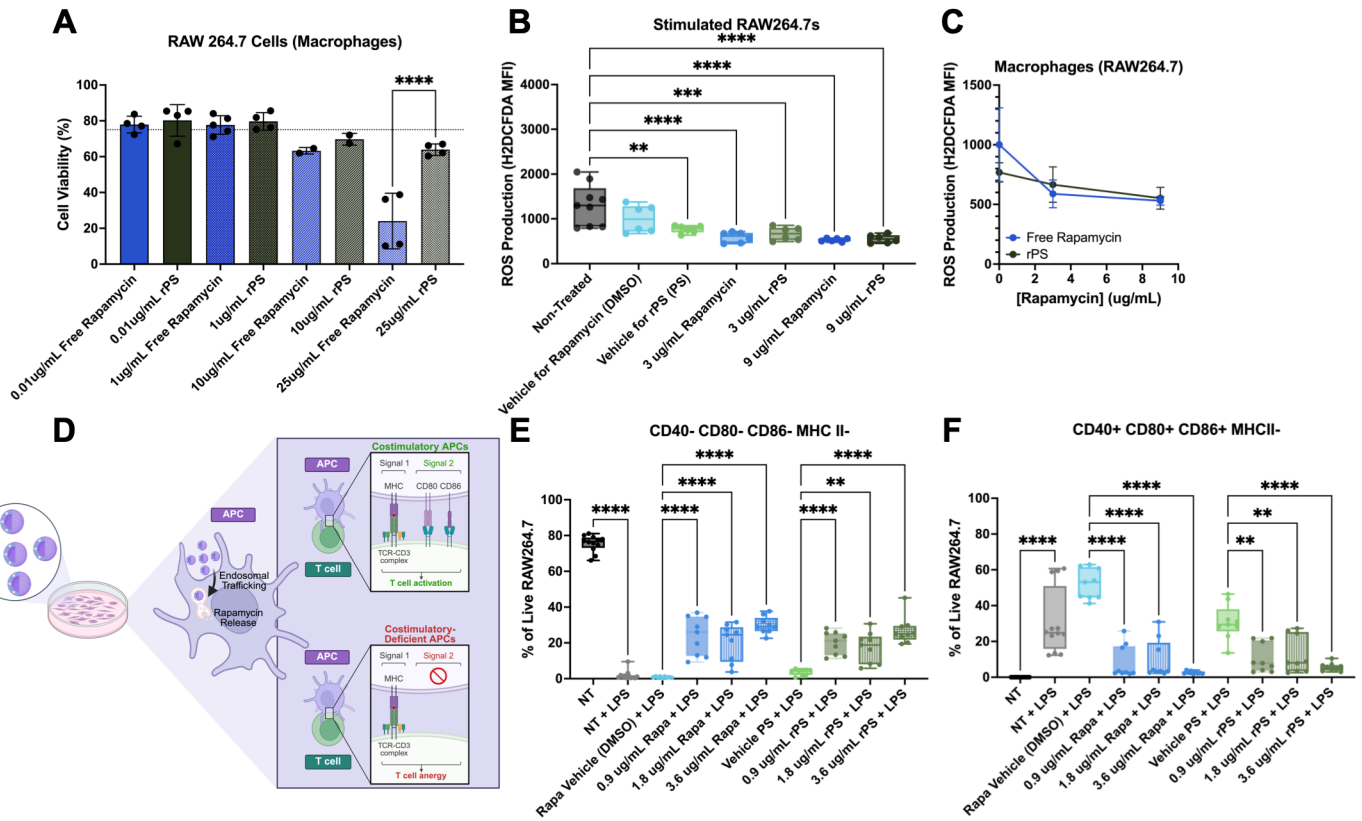

### Supplementary Fig. 9

PS platform modulates APC activation and costimulatory marker expression. (A) rPS treatment supported macrophage (RAW264.7) viability in comparison to free drug. (B) Following LPS stimulation, vehicle PS and all tested formulations of rPS decreased ROS production in macrophages. (C) Increasing the amount of rapamycin in a formulation lowered ROS production. (D) Stimulated macrophages were treated and assessed for costimulatory marker expression. (E) rPS treatment increased the frequency of costimulation-deficient macrophages. (F) rPS treatment decreased the frequency of costimulatory macrophages. Statistical analysis shown within treatment groups. All statistical analyses completed using One-Way ANOVA followed by Tukey's multiple-comparisons test. \*\*P < 0.01, \*\*\*\*P < 0.0001.

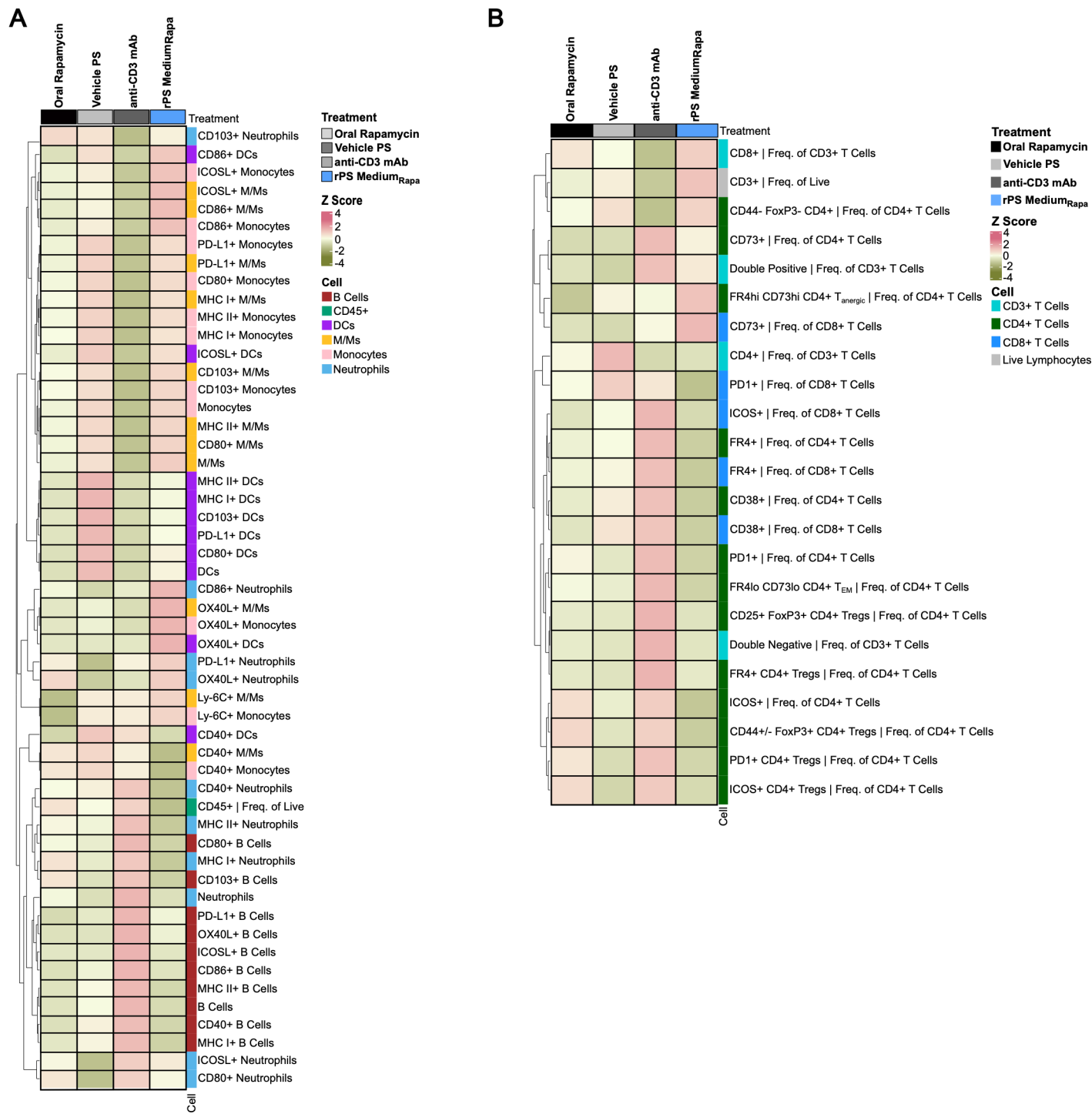

### Supplementary Fig. 10

rPS treatment induces costimulatory-deficient APCs to maintain anergic CD4<sup>+</sup> T cells in vivo. Flow cytometry analysis was performed on the blood collected from non-treated NOD mice or mice treated with oral rapamycin (1 mg rapamycin/kg, Q3D, 6 doses), subcutaneous polymersomes (Q3D, 6 doses), intravenous anti-CD3 mAb (5 ug, QD, 5 doses), or rPS Medium<sub>Rapa</sub> (1 mg rapamycin/kg, Q3D, 6 doses), starting at 8 weeks of age ($n = 4-5$). *A* and *B*: Relative frequency of CD45<sup>+</sup> (*A*) and CD3<sup>+</sup> (*B*) cells shown in heatmap (vs non-treated).

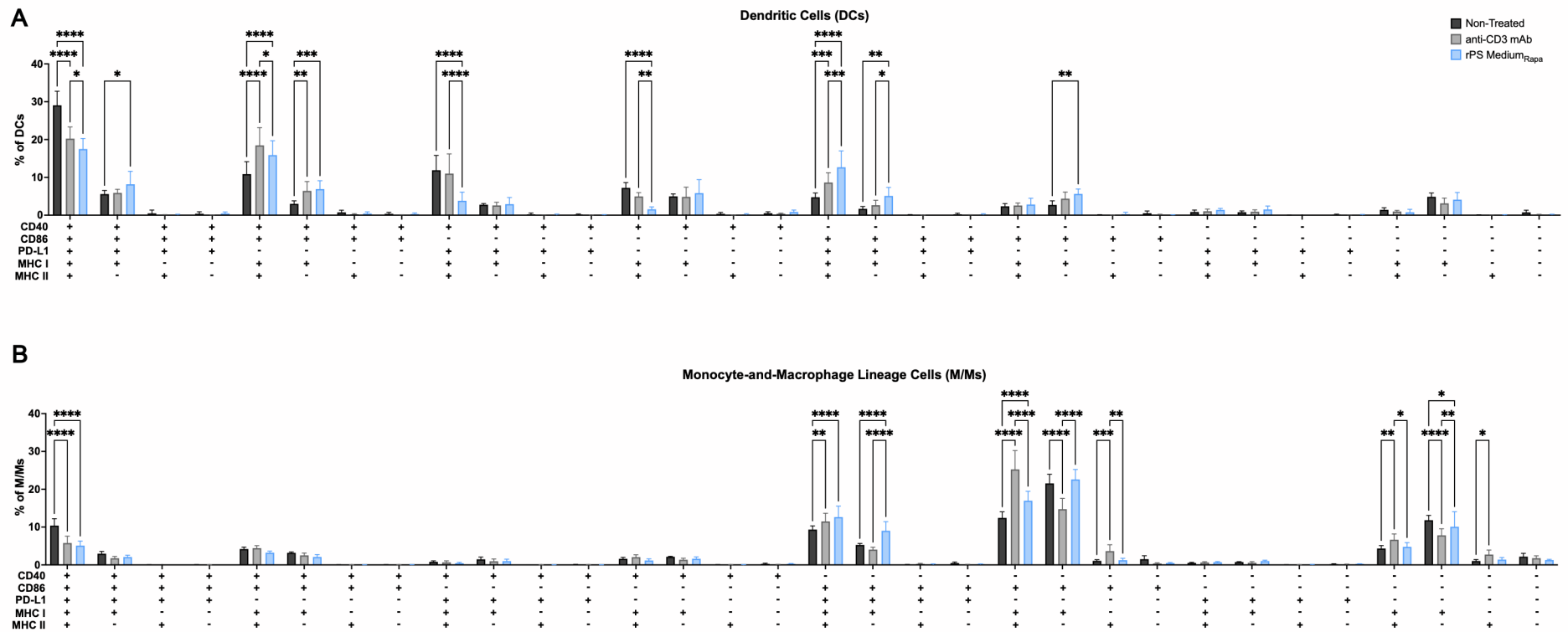

#### Supplementary Fig. 11

rPS treatment induces costimulatory-deficient APCs in vivo. Flow cytometry analysis was performed on the A/B LN collected from non-treated NOD mice or NOD mice treated with oral rapamycin (1 mg rapamycin/kg, Q3D, 6 doses), SC polymersomes (Q3D, 6 doses), IV anti-CD3 mAb (5 ug, QD, 5 doses), or SC rPS Medium<sub>Rapa</sub> (1 mg rapamycin/kg, Q3D, 6 doses), starting at 8 weeks of age ($n = 4-5$). *A*: Relative frequency of each costimulatory combination of DCs. *B*: Relative frequency of each costimulatory combination of M/Ms. Statistical analyses completed using one-way ANOVAs followed by Tukey's multiple-comparisons test. \*$P < 0.05$, \*\*$P < 0.01$, \*\*\*$P < 0.001$, \*\*\*\*$P < 0.0001$.

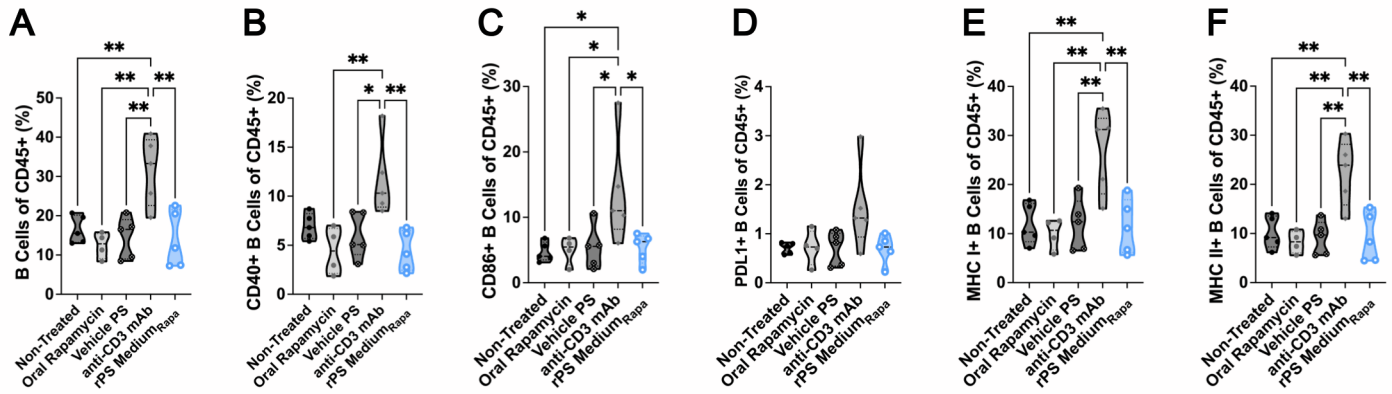

**Supplementary Fig. 12**

rPS SC treatment induces costimulatory-deficient B cells in vivo. Flow cytometry analysis was performed on the A/B LN collected from non-treated NOD mice or mice treated with oral rapamycin (1 mg rapamycin/kg, Q3D, 6 doses), SC polymersomes (Q3D, 6 doses), IV anti-CD3 mAb (5 ug, QD, 5 doses), or SC rPS Medium<sub>Rapa</sub> (1 mg rapamycin/kg, Q3D, 6 doses), starting at 8 weeks of age ($n = 4-5$). *A*: Frequency of B Cells in the A/B LN. *B-F* Frequency of CD40+ (*B*), CD86+ (*C*), PD-L1+ (*D*), MHC I+ (*E*), MHC II+ (*F*) B Cells of CD45+ cells in the A/B LN. Statistical analyses completed using one-way ANOVAs followed by Tukey's multiple-comparisons test. \*$P < 0.05$, \*\*$P < 0.01$.

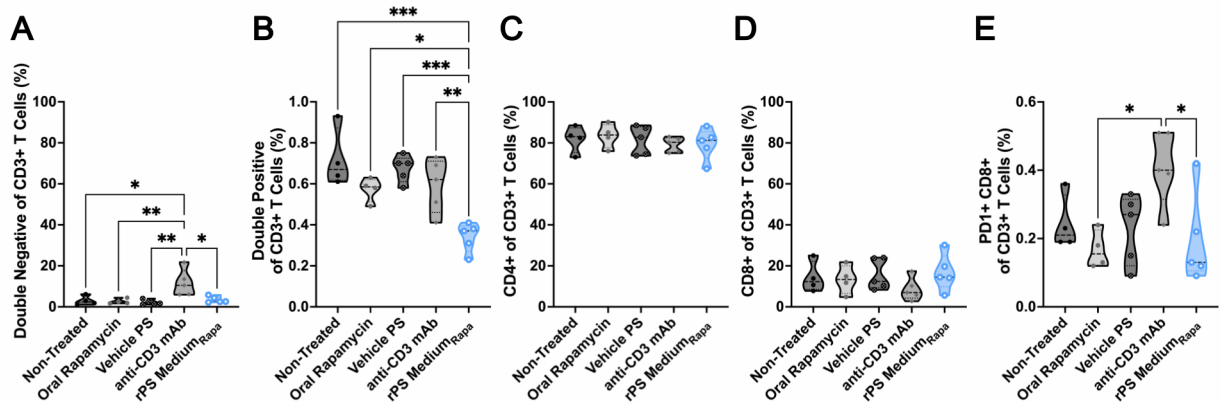

#### Supplementary Fig. 13

rPS treatment avoids lymphodepletion and CD8+ T cell exhaustion in vivo. Flow cytometry analysis was performed on the A/B LN collected from non-treated NOD mice or mice treated with oral rapamycin (1 mg rapamycin/kg, Q3D, 6 doses), SC polymersomes (Q3D, 6 doses), IV anti-CD3 mAb (5 ug, QD, 5 doses), or SC rPS Medium<sub>Rapa</sub> (1 mg rapamycin/kg, Q3D, 6 doses), starting at 8 weeks of age ($n = 4-5$). *A-D*: Overview of major T cell subsets in A/B LN post-treatment. *E*: PD1+ CD8+ T cells of CD3+ T cell in the A/B LN. Statistical analyses completed using one-way ANOVAs followed by Tukey's multiple-comparisons test. \*$P < 0.05$, \*\*$P < 0.01$, \*\*\*$P < 0.001$.

### Supplementary Data Collection Settings and Methods

#### Small angle X-ray scatterings (SAXS).

Data was collected using a set of triple phosphor-coupled CCD area detectors (1) and with the X-ray energy of 17 keV, spanning a $q$-range of 0.003 to 4 Å<sup>-1</sup>, where $q$, the magnitude of the reciprocal space vector, is defined as $q = 4\pi\sin\theta/\lambda$, with $\theta$ denoting half of the total scattering angle and $\lambda$ denoting the X-ray wavelength. A home-built U-shaped sample cell based on laser-cut acrylic plate as spacers and a pair of 50 µm thick glass sheets (G-leaf, Nippon Electric Glass) as the window material was used (https://github.com/stupplab/SAXScells). The scattering contribution of the window materials were first measured by obtaining the scattering patterns of the empty cells, then filling the cells with buffers and samples and measuring the same spots again. The 2D scattering patterns were azimuthally integrated to yield the intensity vs $q$ plots. First the empty cell contribution was subtracted from the buffer and sample patterns, and the buffer contribution was subsequently subtracted from the sample scattering pattern with a scaling factor determined by the algorithm described in (2).

#### PS Biodistribution

To assess the organ-level biodistribution of PS, 8-week-old female NOD mice (n = 3) were subcutaneously injected with 1mg/kg of either free DiR dye or DiR + rapamycin-loaded PS administered in 1, 3, or 6 doses. 24 hours after the respective final dose, the mice were sacrificed, and the liver, heart, kidneys, lungs, spleen, adipose tissue, and pancreas were harvested. In addition, the axial/brachial, inguinal, and pancreatic lymph nodes were collected. These organs and lymph nodes were imaged using the In Vivo Imaging System (IVIS Lumina) with settings for an excitation wavelength of 640 nm, emission wavelength of 680 nm, exposure time of 0.25 seconds, and an aperture of f/2. The images were subsequently analyzed using the IVIS Lumina Living Image Software (v.4.5.5, PerkinElmer).

#### Reactive Oxygen Species (ROS) in vitro Screening Assay

Macrophages (RAW264.7) were cultured according to the supplier's instructions (ATCC). The cell line was seeded at 500,000 cells/well. Cells were incubated for 24 hours with the following treatments: non-treated, free rapamycin in 0.9% DMSO, rPS, vehicle PS, and 0.9% DMSO (N = 9). All nanocarrier formulations matched for polymer concentration (6.7 mg/mL), and the free rapamycin controls were matched to the rapamycin dose encapsulated within the corresponding nanocarrier formulation. The DMSO control was included as the vehicle for free rapamycin. Prior to stimulation and ROS analysis, an Alamar blue assay was performed to confirm cell viability after 24 hours of exposure to each treatment.

Following the 24-hour treatment period, macrophages were stimulated with 1 µg/mL lipopolysaccharide (LPS) for 24 hours, then incubated with H2DCFDA for 25 minutes before fluorescent measurement to assess ROS production.

#### Costimulatory Marker in vitro Flow Cytometry Study

Macrophages (RAW264.7) were cultured according to the supplier's instructions (ATCC). Cells were seeded at 500,000 cells per well in 6-well plates (N = 9). Cells were incubated for 24 hours with the following treatments: non-treated, free rapamycin in 0.9% DMSO, rPS, vehicle PS, and 0.9% DMSO. All nanocarrier formulations matched for polymer concentration (6.7 mg/mL), and the free rapamycin controls were matched to the rapamycin dose encapsulated within the corresponding nanocarrier formulation (1, 2, and 4 µM) and added in equal volume to the appropriate well. The DMSO control was included as the vehicle for free rapamycin..

Following the 24-hour treatment period, macrophages were washed and stimulated with 1 µg/mL LPS for 24 hours. The stimulation media was removed, and the cells were washed then

resuspended in cell staining buffer (BioLegend) for surface staining. The cells were transferred to a 96-well plate and spun down at 400 g for 5 minutes at 4°C. resuspended in a cocktail of LIVE/DEAD Blue (Invitrogen) for viability and anti-mouse CD16/32 (BioLegend) for FcR blocking with cell staining buffer. All cells were incubated at 4°C for 15 minutes in the dark. Cells were washed and spun. The cells were then resuspended in an antibody cocktail consisting of PerCP anti-mouse MHC II (BioLegend), APC anti-mouse CD40 (BioLegend), APC-Fire 750 anti-mouse CD86 (BioLegend), and PE-Dazzle 594 anti-mouse CD80 and incubated with this cocktail for 20 minutes at room temperature in the dark. The cells were washed twice with PBS then resuspended in cell staining buffer for analysis on a 3-laser Aurora flow cytometer (Cytex, Immunotherapy Assessment Core). Spectral unmixing was performed via SpectroFlo (Cytex) and analysis was completed using the FlowJo software.

**Supplementary Tables**

**Supplementary Table 1.** Flow cytometry markers for the APC panel.

| Marker | Fluorophore | Vendor | Clone | Catalog # |
| --- | --- | --- | --- | --- |
| PD-L1 | Spark PLUS UV395 | BioLegend | 10F.9G2 | 124353 |
| Viability | LIVE DEAD Blue | Invitrogen | - | L34966A |
| CD103 | BUV563 | BD | M290 | 741739 |
| MHC I | Spark Violet 423 | BioLegend | H-2Kd/H-2Dd 34-1-2S | 114721 |
| CD80 | BV480 | Invitrogen | 16-10A1 | 414-0801-82 |
| CD11b | BV650 | Biolegend | M1/70 | 101239 |
| CD19 | BV711 | BioLegend | 6D5 | 115555 |
| CD40 | BV785 | BioLegend | 3/23 | 124645 |
| OX40L | BB515 | BD | RM134L | 565214 |
| F4-80 | Spark Blue 550 | BioLegend | BM8 | 123169 |
| CD11c | PerCP-Cy5.5 | BioLegend | N418 | 117327 |
| ICOSL | PE | BioLegend | HK5.3 | 107405 |
| Ly-6C | PE-Dazzle 594 | BioLegend | HK1.4 | 128044 |
| CD86 | PE-Cy7 | Biolegend | GL-1 | 105013 |
| CD45 | PE-Fire 810 | BioLegend | 30-F11 | 103189 |
| MHC II H-2g7 | APC | Invitrogen | AMS-32.1 | 17-5323-82 |
| Ly-6G | APC-Fire810 | BioLegend | 1A8 | 127670 |

**Supplementary Table 2.** Flow cytometry markers for the T cell panel.

| Marker | Fluorophore | Vendor | Clone | Catalog # |
| --- | --- | --- | --- | --- |
| CD3 | BUV395 | BD | 145-2C11 | 563565 |
| Viability | LIVE DEAD Blue | Invitrogen | - | L34966A |
| CD4 | BUV496 | BD | GK1.5 | 612952 |
| CD8 | BUV805 | BD | 53-6.7 | 612898 |
| CD38 | Pacific Blue | BioLegend | 90 | 102719 |
| CD73 | BV605 | BioLegend | TY/11.8 | 127215 |
| ICOS | BV650 | BioLegend | C398.4A | 313550 |
| PD1 | BV711 | BioLegend | 29F.1A12 | 135231 |
| TIGIT | BV786 | BD | 1G9 | 744215 |
| FR4 | PE-Cy7 | BioLegend | 12A5 | 125012 |
| Lag3 | APC | BioLegend | C9B7W | 125209 |
| Foxp3 | AF700 | BioLegend | MF-14 | 126422 |
| CD44 | APC-Fire750 | BioLegend | NIM-R8 | 156004 |
| CD25 | APC-Fire810 | BioLegend | PC61 | 102076 |

**Supplementary Table 3.** B cell subsets overview in A/B LN.

| B Cell Subset | Treatment | Mean Frequency of Cell Type+ of CD45+ Cells | Standard Deviation of Frequency of Cell Type+ of CD45+ Cells | Mean Count of Cell Type+ of CD45+ cells | Standard Deviation of Count of Cell Type+ of CD45+ Cells | Mean Count of CD45+ Cells | Standard Deviation of Count of CD45+ Cells |
| --- | --- | --- | --- | --- | --- | --- | --- |
| CD40+ B Cells | Non-Treated | 6.93% | 1.327 | 37158.4 | 4646.5 | 87524 | 6714 |
|  | Vehicle PS | 5.98% | 2.331 | 30081.8 | 20755.1 | 63894 | 36260 |
|  | rPS Medium<sub>Rapa</sub> | 4.40% | 2.059 | 8858.2 | 5509.9 | 27948 | 18890 |
|  | Oral Rapamycin | 4.42% | 2.417 | 17501.5 | 12934.1 | 46783 | 22472 |
|  | anti-CD3 mAb | 11.73% | 3.902 | 25215.8 | 14805.1 | 61839 | 24717 |
| CD80+ B Cells | Non-Treated | 0.48% | 0.088 | 2585.8 | 386.34 | 87524 | 6714 |
|  | Vehicle PS | 0.67% | 0.414 | 2723.6 | 2160.02 | 63894 | 36260 |
|  | rPS Medium<sub>Rapa</sub> | 0.96% | 0.417 | 1673 | 628.12 | 27948 | 18890 |
|  | Oral Rapamycin | 0.99% | 0.557 | 3349.25 | 1509.69 | 46783 | 22472 |
|  | anti-CD3 mAb | 1.56% | 1.297 | 2588.6 | 1095.61 | 61839 | 24717 |
| CD103+ B Cells | Non-Treated | 1.98% | 1.59 | 10834.2 | 9504.2 | 87524 | 6714 |
|  | Vehicle PS | 3.71% | 4.051 | 20681 | 24950.7 | 63894 | 36260 |
|  | rPS Medium<sub>Rapa</sub> | 3.47% | 3.013 | 5592.2 | 4090 | 27948 | 18890 |
|  | Oral Rapamycin | 4.37% | 3.852 | 15783.5 | 13583.6 | 46783 | 22472 |
|  | anti-CD3 mAb | 8.31% | 12.087 | 11700.8 | 11443.4 | 61839 | 24717 |
| CD86+ B Cells | Non-Treated | 4.69% | 1.495 | 24526.6 | 3263.8 | 87524 | 6714 |
|  | Vehicle PS | 5.39% | 3.308 | 20812.8 | 11374.3 | 63894 | 36260 |
|  | rPS Medium<sub>Rapa</sub> | 5.31% | 2.274 | 9976 | 5466.7 | 27948 | 18890 |
|  | Oral Rapamycin | 4.94% | 2.102 | 17491 | 7373.8 | 46783 | 22472 |
|  | anti-CD3 mAb | 13.91% | 8.197 | 23916 | 5314.4 | 61839 | 24717 |
| MHC I+ B Cells | Non-Treated | 11.85% | 4.144 | 61766.2 | 11702 | 87524 | 6714 |
|  | Vehicle PS | 11.80% | 5.313 | 49671.2 | 27446.1 | 63894 | 36260 |
|  | rPS Medium<sub>Rapa</sub> | 11.44% | 5.558 | 22353 | 13571.3 | 27948 | 18890 |
|  | Oral Rapamycin | 9.93% | 3.165 | 36935 | 17824.6 | 46783 | 22472 |
|  | anti-CD3 mAb | 26.88% | 8.463 | 51497.4 | 19083.3 | 61839 | 24717 |
| MHC II+ B Cells | Non-Treated | 10.15% | 3.314 | 53252 | 10416 | 87524 | 6714 |
|  | Vehicle PS | 9.13% | 3.441 | 39300 | 21249.8 | 63894 | 36260 |
|  | rPS Medium<sub>Rapa</sub> | 9.21% | 4.982 | 18375.8 | 12506.3 | 27948 | 18890 |
|  | Oral Rapamycin | 8.24% | 2.262 | 31106 | 15757.8 | 46783 | 22472 |
|  | anti-CD3 mAb | 22.36% | 6.716 | 43353 | 16335.6 | 61839 | 24717 |
| ICOSL+ B Cells | Non-Treated | 1.02% | 0.267 | 5407.6 | 629 | 87524 | 6714 |
|  | Vehicle PS | 1.50% | 1.156 | 5444 | 3691.7 | 63894 | 36260 |
|  | rPS Medium<sub>Rapa</sub> | 1.95% | 0.798 | 3526.8 | 1628.7 | 27948 | 18890 |
|  | Oral Rapamycin | 1.84% | 0.963 | 6292.5 | 2636.7 | 46783 | 22472 |
|  | anti-CD3 mAb | 4.37% | 4.579 | 6772.8 | 3557.9 | 61839 | 24717 |
| OX40L+ B Cells | Non-Treated | 0.19% | 0.025 | 1062.4 | 175.8 | 87524 | 6714 |
|  | Vehicle PS | 0.22% | 0.111 | 1050 | 825.1 | 63894 | 36260 |
|  | rPS Medium<sub>Rapa</sub> | 0.36% | 0.175 | 636 | 208.8 | 27948 | 18890 |
|  | Oral Rapamycin | 0.36% | 0.311 | 1214.3 | 928.8 | 46783 | 22472 |
|  | anti-CD3 mAb | 0.59% | 0.578 | 896.6 | 437 | 61839 | 24717 |
| PD-L1+ B Cells | Non-Treated | 0.71% | 0.111 | 3867.2 | 792 | 87524 | 6714 |
|  | Vehicle PS | 0.70% | 0.345 | 3301.4 | 2511.6 | 63894 | 36260 |
|  | rPS Medium<sub>Rapa</sub> | 0.69% | 0.297 | 1209 | 419.8 | 27948 | 18890 |
|  | Oral Rapamycin | 0.72% | 0.364 | 2579.3 | 1366.5 | 46783 | 22472 |
|  | anti-CD3 mAb | 1.54% | 0.879 | 2677.2 | 786.9 | 61839 | 24717 |

| DC Subset | Treatment | Mean Frequency of Cell Type+ of CD45+ Cells | Standard Deviation of Frequency of Cell Type+ of CD45+ Cells | Mean Count of Cell Type+ of CD45+ cells | Standard Deviation of Count of Cell Type+ of CD45+ Cells | Mean Count of CD45+ Cells | Standard Deviation of Count of CD45+ Cells |
| --- | --- | --- | --- | --- | --- | --- | --- |
| CD40+ DCs | Non-Treated | 0.46% | 13.10% | 2571 | 1045.4 | 3217 | 1118 |
|  | Vehicle PS | 0.34% | 10.30% | 1567.4 | 940.2 | 1936 | 1086 |
|  | rPS Medium<sub>Rapa</sub> | 0.18% | 6.10% | 361.8 | 185.9 | 569 | 313 |
|  | Oral Rapamycin | 0.26% | 12.50% | 963.5 | 536.8 | 1340 | 640 |
|  | anti-CD3 mAb | 0.50% | 24.30% | 1022.4 | 715.3 | 1329 | 840 |
| CD80+ DCs | Non-Treated | 0.03% | 0.40% | 158.2 | 48.92 | 3217 | 1118 |
|  | Vehicle PS | 0.02% | 1.00% | 80.6 | 38.64 | 1936 | 1086 |
|  | rPS Medium<sub>Rapa</sub> | 0.03% | 1.30% | 51.6 | 37.13 | 569 | 313 |
|  | Oral Rapamycin | 0.03% | 2.10% | 98 | 59.77 | 1080 | 803 |
|  | anti-CD3 mAb | 0.04% | 1.60% | 71.4 | 30.54 | 1329 | 840 |
| CD103+ DCs | Non-Treated | 0.40% | 14.80% | 2227.6 | 1067.3 | 3217 | 1118 |
|  | Vehicle PS | 0.30% | 8.20% | 1458.6 | 901.6 | 1936 | 1086 |
|  | rPS Medium<sub>Rapa</sub> | 0.18% | 7.00% | 354 | 176 | 569 | 313 |
|  | Oral Rapamycin | 0.28% | 15.40% | 1017.5 | 607.8 | 1080 | 803 |
|  | anti-CD3 mAb | 0.41% | 16.60% | 797.6 | 402.5 | 1329 | 840 |
| CD86+ DCs | Non-Treated | 0.36% | 7.30% | 2017.2 | 675.4 | 3217 | 1118 |
|  | Vehicle PS | 0.29% | 11.90% | 1247.2 | 620.2 | 1936 | 1086 |
|  | rPS Medium<sub>Rapa</sub> | 0.22% | 9.00% | 426.2 | 228.2 | 569 | 313 |
|  | Oral Rapamycin | 0.26% | 12.10% | 943 | 455.2 | 1080 | 803 |
|  | anti-CD3 mAb | 0.45% | 18.00% | 880.4 | 470.4 | 1329 | 840 |
| MHC I+ DCs | Non-Treated | 0.55% | 12.40% | 3041.4 | 1043 | 3217 | 1118 |
|  | Vehicle PS | 0.41% | 12.40% | 1870.4 | 1034.2 | 1936 | 1086 |
|  | rPS Medium<sub>Rapa</sub> | 0.28% | 11.80% | 551.8 | 303.8 | 569 | 313 |
|  | Oral Rapamycin | 0.35% | 13.80% | 1303.8 | 626.6 | 1080 | 803 |
|  | anti-CD3 mAb | 0.64% | 27.10% | 1311.8 | 832.1 | 1329 | 840 |
| MHC II+ DCs | Non-Treated | 0.41% | 9.40% | 2287 | 855 | 3217 | 1118 |
|  | Vehicle PS | 0.28% | 8.40% | 1255 | 720.3 | 1936 | 1086 |
|  | rPS Medium<sub>Rapa</sub> | 0.17% | 8.70% | 340.8 | 213.7 | 569 | 313 |
|  | Oral Rapamycin | 0.24% | 11.20% | 866 | 418.2 | 1080 | 803 |
|  | anti-CD3 mAb | 0.44% | 17.60% | 884.4 | 511.2 | 1329 | 840 |
| ICOSL+ DCs | Non-Treated | 0.01% | 0.30% | 74 | 24.1 | 3217 | 1118 |
|  | Vehicle PS | 0.02% | 1.80% | 65.8 | 44.2 | 1936 | 1086 |
|  | rPS Medium<sub>Rapa</sub> | 0.02% | 1.50% | 39.2 | 18.9 | 569 | 313 |
|  | Oral Rapamycin | 0.03% | 2.90% | 89.8 | 87.8 | 1080 | 803 |
|  | anti-CD3 mAb | 0.04% | 4.90% | 65.8 | 41 | 1329 | 840 |
| OX40L+<br>DCs | Non-Treated | 0.04% | 0.90% | 199.8 | 84.4 | 3217 | 1118 |
|  | Vehicle PS | 0.03% | 1.60% | 111 | 56.9 | 1936 | 1086 |
|  | rPS Medium<sub>Rapa</sub> | 0.03% | 1.20% | 53.4 | 34.4 | 569 | 313 |
|  | Oral Rapamycin | 0.04% | 3.80% | 134 | 110.5 | 1080 | 803 |
|  | anti-CD3 mAb | 0.04% | 3.10% | 70.8 | 29.3 | 1329 | 840 |
| PD-L1+<br>DCs | Non-Treated | 0.35% | 11.30% | 1959.4 | 872 | 3217 | 1118 |
|  | Vehicle PS | 0.23% | 6.20% | 1106 | 681.6 | 1936 | 1086 |
|  | rPS Medium<sub>Rapa</sub> | 0.15% | 5.70% | 297.6 | 149.5 | 569 | 313 |
|  | Oral Rapamycin | 0.21% | 10.30% | 764.8 | 401.9 | 1080 | 803 |
|  | anti-CD3 mAb | 0.35% | 15.00% | 712.2 | 464.7 | 1329 | 840 |

| M/M Subsets | Treatment | Mean Frequency of Cell Type+ of CD45+ Cells | Standard Deviation of Frequency of Cell Type+ of CD45+ Cells | Mean Count of Cell Type+ of CD45+ cells | Standard Deviation of Count of Cell Type+ of CD45+ Cells | Mean Count of CD45+ Cells | Standard Deviation of Count of CD45+ Cells |
| --- | --- | --- | --- | --- | --- | --- | --- |
| CD40+ M/Ms | Non-Treated | 0.48% | 0.08 | 2680 | 965.9 | 9810 | 3345 |
|  | Vehicle PS | 0.29% | 0.06 | 1398.4 | 818.8 | 5231 | 2487 |
|  | rPS Medium<sub>Rapa</sub> | 0.29% | 0.16 | 618.4 | 452.5 | 3448 | 2388 |
|  | Oral Rapamycin | 0.34% | 0.17 | 1187.8 | 485.9 | 6244 | 2251 |
|  | anti-CD3 mAb | 0.82% | 0.32 | 1530.6 | 583.2 | 7637 | 1696 |
| CD80+ M/Ms | Non-Treated | 0.30% | 0.03 | 1653.2 | 443.69 | 9810 | 3345 |
|  | Vehicle PS | 0.24% | 0.08 | 1040 | 583.77 | 5231 | 2487 |
|  | rPS Medium<sub>Rapa</sub> | 0.41% | 0.16 | 812.8 | 481.03 | 3448 | 2388 |
|  | Oral Rapamycin | 0.46% | 0.31 | 1498.5 | 796.97 | 6244 | 2251 |
|  | anti-CD3 mAb | 0.84% | 0.61 | 1401.8 | 499.87 | 7637 | 1696 |
| CD103+ M/Ms | Non-Treated | 0.42% | 0.09 | 2335.8 | 802.3 | 9810 | 3345 |
|  | Vehicle PS | 0.38% | 0.21 | 1937 | 1540 | 5231 | 2487 |
|  | rPS Medium<sub>Rapa</sub> | 0.53% | 0.28 | 985.6 | 603 | 3448 | 2388 |
|  | Oral Rapamycin | 0.85% | 0.75 | 2775 | 2209.5 | 6244 | 2251 |
|  | anti-CD3 mAb | 1.47% | 1.83 | 2128.2 | 1584.1 | 7637 | 1696 |
| CD86+ M/Ms | Non-Treated | 1.28% | 0.23 | 7093.4 | 2271.1 | 9810 | 3345 |
|  | Vehicle PS | 0.91% | 0.24 | 3939.8 | 1830.8 | 5231 | 2487 |
|  | rPS Medium<sub>Rapa</sub> | 1.24% | 0.61 | 2533.4 | 1675.6 | 3448 | 2388 |
|  | Oral Rapamycin | 1.48% | 0.91 | 4864.3 | 2187.2 | 6244 | 2251 |
|  | anti-CD3 mAb | 3.33% | 1.85 | 5681.8 | 1325.3 | 7637 | 1696 |
| MHC I+ M/Ms | Non-Treated | 1.64% | 0.3 | 9067.2 | 2911.8 | 9810 | 3345 |
|  | Vehicle PS | 1.12% | 0.3 | 4881.4 | 2304 | 5231 | 2487 |
|  | rPS Medium<sub>Rapa</sub> | 1.58% | 0.9 | 3258.4 | 2253.8 | 3448 | 2388 |
|  | Oral Rapamycin | 1.78% | 1 | 5908.3 | 2292.9 | 6244 | 2251 |
|  | anti-CD3 mAb | 3.99% | 2 | 6910.4 | 1461.9 | 7637 | 1696 |
| MHC II+ M/Ms | Non-Treated | 0.80% | 0.12 | 4459 | 1305.5 | 9810 | 3345 |
|  | Vehicle PS | 0.62% | 0.22 | 2572.2 | 1058.5 | 5231 | 2487 |
|  | rPS Medium<sub>Rapa</sub> | 0.78% | 0.4 | 1600 | 1064 | 3448 | 2388 |
|  | Oral Rapamycin | 0.89% | 0.43 | 2998.5 | 817.8 | 6244 | 2251 |
|  | anti-CD3 mAb | 2.79% | 1.41 | 4888 | 1234.1 | 7637 | 1696 |
| ICOSL+ M/Ms | Non-Treated | 0.16% | 0.02 | 884.8 | 247.4 | 9810 | 3345 |
|  | Vehicle PS | 0.19% | 0.11 | 758.6 | 535.7 | 5231 | 2487 |
|  | rPS Medium<sub>Rapa</sub> | 0.34% | 0.17 | 612.2 | 306.7 | 3448 | 2388 |
|  | Oral Rapamycin | 0.42% | 0.34 | 1373.8 | 944.4 | 6244 | 2251 |
|  | anti-CD3 mAb | 0.98% | 1.14 | 1406.8 | 992.5 | 7637 | 1696 |
| OX40L+<br>M/Ms | Non-Treated | 0.70% | 0.23 | 3942.4 | 1893.4 | 9810 | 3345 |
|  | Vehicle PS | 0.41% | 0.11 | 1983.2 | 1162.2 | 5231 | 2487 |
|  | rPS Medium<sub>Rapa</sub> | 0.69% | 0.35 | 1442.8 | 951.6 | 3448 | 2388 |
|  | Oral Rapamycin | 0.91% | 0.75 | 2945 | 2107.2 | 6244 | 2251 |
|  | anti-CD3 mAb | 1.06% | 0.72 | 1676 | 381.1 | 7637 | 1696 |
| PD-L1+<br>M/Ms | Non-Treated | 0.57% | 0.1 | 3188.6 | 1131.5 | 9810 | 3345 |
|  | Vehicle PS | 0.37% | 0.08 | 1774.4 | 1019.9 | 5231 | 2487 |
|  | rPS Medium<sub>Rapa</sub> | 0.52% | 0.23 | 1057.8 | 654.1 | 3448 | 2388 |
|  | Oral Rapamycin | 0.63% | 0.35 | 2117 | 882.5 | 6244 | 2251 |
|  | anti-CD3 mAb | 1.13% | 0.51 | 2042.4 | 565.4 | 7637 | 1696 |
| Ly-6C+<br>M/Ms | Non-Treated | 0.10% | 0.02 | 535 | 196.7 | 9810 | 3345 |
|  | Vehicle PS | 0.10% | 0.06 | 343.6 | 67.1 | 5231 | 2487 |
|  | rPS Medium<sub>Rapa</sub> | 0.12% | 0.03 | 234.4 | 113.7 | 3448 | 2388 |
|  | Oral Rapamycin | 0.15% | 0.1 | 473.5 | 269.2 | 6244 | 2251 |
|  | anti-CD3 mAb | 0.25% | 0.16 | 403.8 | 69.3 | 7637 | 1696 |

Supplementary Table 6. B cell subsets overview in blood.

| B Cell Subset | Treatment | Mean Frequency of Cell Type+ of CD45+ Cells | Standard Deviation of Frequency of Cell Type+ of CD45+ Cells | Mean Count of Cell Type+ of CD45+ cells | Standard Deviation of Count of Cell Type+ of CD45+ Cells | Mean Count of CD45+ Cells | Standard Deviation of Count of CD45+ Cells |
| --- | --- | --- | --- | --- | --- | --- | --- |
| CD40+ B Cells | Non-Treated | 6.93% | 1.327 | 37158.4 | 4646.5 | 87524 | 6714 |
|  | Vehicle PS | 5.98% | 2.331 | 30081.8 | 20755.1 | 63894 | 36260 |
|  | rPS Medium<sub>Rapa</sub> | 4.40% | 2.059 | 8858.2 | 5509.9 | 27948 | 18890 |
|  | Oral Rapamycin | 4.42% | 2.417 | 17501.5 | 12934.1 | 46783 | 22472 |
|  | anti-CD3 mAb | 11.73% | 3.902 | 25215.8 | 14805.1 | 61839 | 24717 |
| CD80+ B Cells | Non-Treated | 0.48% | 0.088 | 2585.8 | 386.34 | 87524 | 6714 |
|  | Vehicle PS | 0.67% | 0.414 | 2723.6 | 2160.02 | 63894 | 36260 |
|  | rPS Medium<sub>Rapa</sub> | 0.96% | 0.417 | 1673 | 628.12 | 27948 | 18890 |
|  | Oral Rapamycin | 0.99% | 0.557 | 3349.25 | 1509.69 | 46783 | 22472 |
|  | anti-CD3 mAb | 1.56% | 1.297 | 2588.6 | 1095.61 | 61839 | 24717 |
| CD103+ B Cells | Non-Treated | 1.98% | 1.59 | 10834.2 | 9504.2 | 87524 | 6714 |
|  | Vehicle PS | 3.71% | 4.051 | 20681 | 24950.7 | 63894 | 36260 |
|  | rPS Medium<sub>Rapa</sub> | 3.47% | 3.013 | 5592.2 | 4090 | 27948 | 18890 |
|  | Oral Rapamycin | 4.37% | 3.852 | 15783.5 | 13583.6 | 46783 | 22472 |
|  | anti-CD3 mAb | 8.31% | 12.087 | 11700.8 | 11443.4 | 61839 | 24717 |
| CD86+ B Cells | Non-Treated | 4.69% | 1.495 | 24526.6 | 3263.8 | 87524 | 6714 |
|  | Vehicle PS | 5.39% | 3.308 | 20812.8 | 11374.3 | 63894 | 36260 |
|  | rPS Medium<sub>Rapa</sub> | 5.31% | 2.274 | 9976 | 5466.7 | 27948 | 18890 |
|  | Oral Rapamycin | 4.94% | 2.102 | 17491 | 7373.8 | 46783 | 22472 |
|  | anti-CD3 mAb | 13.91% | 8.197 | 23916 | 5314.4 | 61839 | 24717 |
| MHC I+ B Cells | Non-Treated | 11.85% | 4.144 | 61766.2 | 11702 | 87524 | 6714 |
|  | Vehicle PS | 11.80% | 5.313 | 49671.2 | 27446.1 | 63894 | 36260 |
|  | rPS Medium<sub>Rapa</sub> | 11.44% | 5.558 | 22353 | 13571.3 | 27948 | 18890 |
|  | Oral Rapamycin | 9.93% | 3.165 | 36935 | 17824.6 | 46783 | 22472 |
|  | anti-CD3 mAb | 26.88% | 8.463 | 51497.4 | 19083.3 | 61839 | 24717 |
| MHC II+ B Cells | Non-Treated | 10.15% | 3.314 | 53252 | 10416 | 87524 | 6714 |
|  | Vehicle PS | 9.13% | 3.441 | 39300 | 21249.8 | 63894 | 36260 |
|  | rPS Medium<sub>Rapa</sub> | 9.21% | 4.982 | 18375.8 | 12506.3 | 27948 | 18890 |
|  | Oral Rapamycin | 8.24% | 2.262 | 31106 | 15757.8 | 46783 | 22472 |
|  | anti-CD3 mAb | 22.36% | 6.716 | 43353 | 16335.6 | 61839 | 24717 |
| ICOSL+ B Cells | Non-Treated | 1.02% | 0.267 | 5407.6 | 629 | 87524 | 6714 |
|  | Vehicle PS | 1.50% | 1.156 | 5444 | 3691.7 | 63894 | 36260 |
|  | rPS Medium<sub>Rapa</sub> | 1.95% | 0.798 | 3526.8 | 1628.7 | 27948 | 18890 |
|  | Oral Rapamycin | 1.84% | 0.963 | 6292.5 | 2636.7 | 46783 | 22472 |
|  | anti-CD3 mAb | 4.37% | 4.579 | 6772.8 | 3557.9 | 61839 | 24717 |
| OX40L+ B Cells | Non-Treated | 0.19% | 0.025 | 1062.4 | 175.8 | 87524 | 6714 |
|  | Vehicle PS | 0.22% | 0.111 | 1050 | 825.1 | 63894 | 36260 |
|  | rPS Medium<sub>Rapa</sub> | 0.36% | 0.175 | 636 | 208.8 | 27948 | 18890 |
|  | Oral Rapamycin | 0.36% | 0.311 | 1214.3 | 928.8 | 46783 | 22472 |
|  | anti-CD3 mAb | 0.59% | 0.578 | 896.6 | 437 | 61839 | 24717 |
| PD-L1+ B Cells | Non-Treated | 0.71% | 0.111 | 3867.2 | 792 | 87524 | 6714 |
|  | Vehicle PS | 0.70% | 0.345 | 3301.4 | 2511.6 | 63894 | 36260 |
|  | rPS Medium<sub>Rapa</sub> | 0.69% | 0.297 | 1209 | 419.8 | 27948 | 18890 |
|  | Oral Rapamycin | 0.72% | 0.364 | 2579.3 | 1366.5 | 46783 | 22472 |
|  | anti-CD3 mAb | 1.54% | 0.879 | 2677.2 | 786.9 | 61839 | 24717 |

**Supplementary Table 7.** DC subsets overview in blood.

| DC Subset | Treatment | Mean Frequency of Cell Type+ of CD45+ Cells | Standard Deviation of Frequency of Cell Type+ of CD45+ Cells | Mean Count of Cell Type+ of CD45+ cells | Standard Deviation of Count of Cell Type+ of CD45+ Cells | Mean Count of CD45+ Cells | Standard Deviation of Count of CD45+ Cells |
| --- | --- | --- | --- | --- | --- | --- | --- |
| CD40+ DCs | Non-Treated | 0.74% | 0.48 | 101.6 | 118.7 | 188 | 226 |
|  | Vehicle PS | 0.65% | 0.145 | 89.2 | 55.1 | 243 | 222 |
|  | rPS Medium<sub>Rapa</sub> | 0.38% | 0.102 | 14.8 | 13.7 | 59 | 51 |
|  | Oral Rapamycin | 0.40% | 0.094 | 88 | 104.7 | 168 | 206 |
|  | anti-CD3 mAb | 0.57% | 0.305 | 75.4 | 85.9 | 111 | 104 |
| CD80+ DCs | Non-Treated | 1.32% | 0.737 | 172 | 220.35 | 188 | 226 |
|  | Vehicle PS | 1.51% | 0.922 | 234.6 | 220.77 | 243 | 222 |
|  | rPS Medium<sub>Rapa</sub> | 1.27% | 0.61 | 55.75 | 48.79 | 59 | 51 |
|  | Oral Rapamycin | 0.82% | 0.388 | 159.8 | 200.47 | 168 | 206 |
|  | anti-CD3 mAb | 0.86% | 0.403 | 99.6 | 87.89 | 111 | 104 |
| CD103+ DCs | Non-Treated | 1.30% | 0.659 | 172.8 | 210.7 | 188 | 226 |
|  | Vehicle PS | 1.43% | 0.868 | 223.4 | 209.4 | 243 | 222 |
|  | rPS Medium<sub>Rapa</sub> | 1.11% | 0.578 | 51.5 | 46.9 | 59 | 51 |
|  | Oral Rapamycin | 0.75% | 0.324 | 150.6 | 191.6 | 168 | 206 |
|  | anti-CD3 mAb | 0.74% | 0.327 | 87.8 | 83.1 | 111 | 104 |
| CD86+ DCs | Non-Treated | 0.67% | 0.642 | 70.6 | 72.9 | 188 | 226 |
|  | Vehicle PS | 0.55% | 0.493 | 87 | 104.4 | 243 | 222 |
|  | rPS Medium<sub>Rapa</sub> | 0.71% | 0.58 | 31.5 | 32.3 | 59 | 51 |
|  | Oral Rapamycin | 0.23% | 0.16 | 35.8 | 32.5 | 168 | 206 |
|  | anti-CD3 mAb | 0.20% | 0.173 | 17.4 | 14.5 | 111 | 104 |
| MHC I+ DCs | Non-Treated | 1.46% | 0.738 | 185.8 | 223.7 | 188 | 226 |
|  | Vehicle PS | 1.54% | 0.93 | 239.2 | 224.1 | 243 | 222 |
|  | rPS Medium<sub>Rapa</sub> | 1.27% | 0.604 | 57 | 50.9 | 59 | 51 |
|  | Oral Rapamycin | 0.85% | 0.352 | 166.6 | 205.2 | 168 | 206 |
|  | anti-CD3 mAb | 0.91% | 0.378 | 110.4 | 103 | 111 | 104 |
| MHC II+ DCs | Non-Treated | 1.44% | 0.746 | 184 | 223.9 | 188 | 226 |
|  | Vehicle PS | 1.56% | 0.913 | 240.6 | 221.5 | 243 | 222 |
|  | rPS Medium<sub>Rapa</sub> | 1.25% | 0.596 | 56.3 | 50 | 59 | 51 |
|  | Oral Rapamycin | 0.86% | 0.377 | 165.6 | 202.9 | 168 | 206 |
|  | anti-CD3 mAb | 0.91% | 0.369 | 110 | 103.5 | 111 | 104 |
| ICOSL+ DCs | Non-Treated | 0.86% | 0.851 | 101.6 | 121.7 | 188 | 226 |
|  | Vehicle PS | 1.00% | 0.915 | 166.6 | 194.2 | 243 | 222 |
|  | rPS Medium<sub>Rapa</sub> | 1.07% | 0.685 | 48 | 44.9 | 59 | 51 |
|  | Oral Rapamycin | 0.52% | 0.468 | 90.2 | 121.3 | 168 | 206 |
|  | anti-CD3 mAb | 0.44% | 0.524 | 42.8 | 46.4 | 111 | 104 |
| OX40L+<br>DCs | Non-Treated | 0.13% | 0.07 | 18.2 | 24.5 | 188 | 226 |
|  | Vehicle PS | 0.08% | 0.057 | 12.6 | 13 | 243 | 222 |
|  | rPS Medium<sub>Rapa</sub> | 0.27% | 0.174 | 14.8 | 17.1 | 59 | 51 |
|  | Oral Rapamycin | 0.08% | 0.03 | 16.2 | 21.8 | 168 | 206 |
|  | anti-CD3 mAb | 0.14% | 0.146 | 15.8 | 13.7 | 111 | 104 |
| PD-L1+<br>DCs | Non-Treated | 1.42% | 0.753 | 178.2 | 213.3 | 188 | 226 |
|  | Vehicle PS | 1.52% | 0.891 | 234.4 | 216.3 | 243 | 222 |
|  | rPS Medium<sub>Rapa</sub> | 1.28% | 0.618 | 56.3 | 49.4 | 59 | 51 |
|  | Oral Rapamycin | 0.81% | 0.36 | 152.8 | 181.2 | 168 | 206 |
|  | anti-CD3 mAb | 0.82% | 0.354 | 96.4 | 88.5 | 111 | 104 |

| M/M Subsets | Treatment | Mean Frequency of Cell Type+ of CD45+ Cells | Standard Deviation of Frequency of Cell Type+ of CD45+ Cells | Mean Count of Cell Type+ of CD45+ cells | Standard Deviation of Count of Cell Type+ of CD45+ T Cells | Mean Count of CD45+ Cells | Standard Deviation of Count of CD45+ Cells |
| --- | --- | --- | --- | --- | --- | --- | --- |
| CD40+ M/Ms | Non-Treated | 0.70% | 0.34 | 92.8 | 139.7 | 345 | 454 |
|  | Vehicle PS | 0.57% | 0.22 | 75.4 | 44.4 | 454 | 432 |
|  | rPS Medium<sub>Rapa</sub> | 0.32% | 0.18 | 17 | 21.3 | 147 | 134 |
|  | Oral Rapamycin | 0.52% | 0.18 | 114.8 | 135.8 | 401 | 434 |
|  | anti-CD3 mAb | 0.46% | 0.3 | 64.8 | 80.3 | 335 | 351 |
| CD80+ M/Ms | Non-Treated | 3.00% | 3.01 | 339.6 | 446 | 345 | 454 |
|  | Vehicle PS | 3.01% | 2.03 | 446.8 | 432.98 | 454 | 432 |
|  | rPS Medium<sub>Rapa</sub> | 3.47% | 2.45 | 141.25 | 132.95 | 147 | 134 |
|  | Oral Rapamycin | 2.42% | 1.31 | 393 | 427.64 | 401 | 434 |
|  | anti-CD3 mAb | 2.11% | 1.3 | 325.6 | 340.33 | 335 | 351 |
| CD103+ M/Ms | Non-Treated | 1.79% | 1.42 | 204 | 257.5 | 345 | 454 |
|  | Vehicle PS | 1.48% | 1.23 | 236.8 | 267.2 | 454 | 432 |
|  | rPS Medium<sub>Rapa</sub> | 1.78% | 1.56 | 80.3 | 84.4 | 147 | 134 |
|  | Oral Rapamycin | 1.24% | 0.61 | 219.8 | 256.5 | 401 | 434 |
|  | anti-CD3 mAb | 0.87% | 0.54 | 118.6 | 116.2 | 335 | 351 |
| CD86+ M/Ms | Non-Treated | 1.13% | 1.92 | 98.2 | 123.3 | 345 | 454 |
|  | Vehicle PS | 0.84% | 0.88 | 124 | 170.5 | 454 | 432 |
|  | rPS Medium<sub>Rapa</sub> | 1.69% | 1.61 | 61.5 | 75 | 147 | 134 |
|  | Oral Rapamycin | 0.51% | 0.5 | 66 | 57.7 | 401 | 434 |
|  | anti-CD3 mAb | 0.24% | 0.24 | 21.4 | 18.8 | 335 | 351 |
| MHC I+ M/Ms | Non-Treated | 2.86% | 2.7 | 332.6 | 441.5 | 345 | 454 |
|  | Vehicle PS | 2.74% | 2 | 421 | 435.3 | 454 | 432 |
|  | rPS Medium<sub>Rapa</sub> | 3.12% | 2.4 | 136.5 | 135.4 | 147 | 134 |
|  | Oral Rapamycin | 2.28% | 1 | 385.8 | 428.3 | 401 | 434 |
|  | anti-CD3 mAb | 2.03% | 1.4 | 318 | 332 | 335 | 351 |
| MHC II+ M/Ms | Non-Treated | 3.00% | 2.98 | 340 | 446.7 | 345 | 454 |
|  | Vehicle PS | 3.04% | 2.01 | 448.4 | 430.8 | 454 | 432 |
|  | rPS Medium<sub>Rapa</sub> | 3.44% | 2.45 | 141.8 | 134.1 | 147 | 134 |
|  | Oral Rapamycin | 2.42% | 1.29 | 396 | 432.8 | 401 | 434 |
|  | anti-CD3 mAb | 2.08% | 1.32 | 322.4 | 337.3 | 335 | 351 |
| ICOSL+ M/Ms | Non-Treated | 2.06% | 2.73 | 205.6 | 235.8 | 345 | 454 |
|  | Vehicle PS | 1.89% | 1.7 | 291 | 348.7 | 454 | 432 |
|  | rPS Medium<sub>Rapa</sub> | 2.98% | 2.53 | 114.3 | 120.9 | 147 | 134 |
|  | Oral Rapamycin | 1.50% | 1.51 | 204.6 | 216.4 | 401 | 434 |
|  | anti-CD3 mAb | 1.35% | 1.45 | 168.4 | 150.8 | 335 | 351 |
| OX40L+<br>M/Ms | Non-Treated | 0.12% | 0.06 | 17.4 | 26.8 | 345 | 454 |
|  | Vehicle PS | 0.13% | 0.15 | 20.2 | 28.9 | 454 | 432 |
|  | rPS Medium<sub>Rapa</sub> | 0.27% | 0.31 | 17.3 | 21.1 | 147 | 134 |
|  | Oral Rapamycin | 0.11% | 0.05 | 23.2 | 28.1 | 401 | 434 |
|  | anti-CD3 mAb | 0.22% | 0.28 | 28.6 | 26.5 | 335 | 351 |
| PD-L1+<br>M/Ms | Non-Treated | 2.62% | 2.9 | 282.8 | 364.4 | 345 | 454 |
|  | Vehicle PS | 2.60% | 1.8 | 382.4 | 375.9 | 454 | 432 |
|  | rPS Medium<sub>Rapa</sub> | 2.80% | 2.43 | 120.8 | 128.6 | 147 | 134 |
|  | Oral Rapamycin | 1.99% | 1.22 | 304.4 | 315.1 | 401 | 434 |
|  | anti-CD3 mAb | 1.74% | 0.98 | 264.6 | 274.3 | 335 | 351 |
| Ly-6C+<br>M/Ms | Non-Treated | 0.05% | 0.04 | 6.6 | 11.5 | 345 | 454 |
|  | Vehicle PS | 0.14% | 0.18 | 16.2 | 15.8 | 454 | 432 |
|  | rPS Medium<sub>Rapa</sub> | 0.08% | 0.08 | 4.5 | 7 | 147 | 134 |
|  | Oral Rapamycin | 0.04% | 0.05 | 3 | 3 | 401 | 434 |
|  | anti-CD3 mAb | 0.17% | 0.16 | 17 | 16.4 | 335 | 351 |

|  |  | Cell Type | Treatment | Mean Frequency of Cell Type+ of CD3+ T cells | Standard Deviation of Frequency of Cell Type+ of CD3+ T cells | Mean Count of Cell Type+ of CD3+ T cells | Standard Deviation of Count of Cell Type+ of CD3+ T cells | Mean Count of CD3+ T cells | Standard Deviation of Count of CD3+ T cells |
| --- | --- | --- | --- | --- | --- | --- | --- | --- | --- |
| Blood | CD8+ CD4- T cells | Non-Treated |  | 29.92% | 3.73% | 4239 | 6122 | 15601 | 23169 |
|  |  | Vehicle PS |  | 29.12% | 2.64% | 5801 | 6339 | 21044 | 24631 |
|  |  | rPS Medium<sub>Rapa</sub> |  | 37.98% | 5.32% | 3535 | 2232 | 9391 | 5596 |
|  |  | Oral Rapamycin |  | 33.30% | 2.76% | 4229 | 4818 | 13393 | 16075 |
|  |  | anti-CD3 mAb |  | 15.91% | 6.68% | 1180 | 492 | 8523 | 3993 |
|  | CD8+ CD4+ (Double Positive) T cells | Non-Treated |  | 0.21% | 0.12% | 20.6 | 28.2 | 15601 | 23169 |
|  |  | Vehicle PS |  | 0.11% | 0.05% | 31.6 | 48.7 | 21044 | 24631 |
|  |  | rPS Medium<sub>Rapa</sub> |  | 0.12% | 0.04% | 10.8 | 6.1 | 9391 | 5596 |
|  |  | Oral Rapamycin |  | 0.11% | 0.03% | 14 | 16 | 13393 | 16075 |
|  |  | anti-CD3 mAb |  | 0.13% | 0.10% | 9.8 | 10.9 | 8523 | 3993 |
|  | CD8- CD4+ T cells | Non-Treated |  | 66.02% | 6.70% | 11080 | 16769 | 15601 | 23169 |
|  |  | Vehicle PS |  | 67.36% | 3.19% | 14640 | 17729 | 21044 | 24631 |
|  |  | rPS Medium<sub>Rapa</sub> |  | 58.50% | 5.87% | 5556 | 3362 | 9391 | 5596 |
|  |  | Oral Rapamycin |  | 61.62% | 4.80% | 8740 | 10970 | 13393 | 16075 |
|  |  | anti-CD3 mAb |  | 57.60% | 15.49% | 4554 | 1844 | 8523 | 3993 |
|  | CD8- CD4- (Double Negative) | Non-Treated |  | 3.87% | 3.16% | 261 | 253 | 15601 | 23169 |
|  |  | Vehicle PS |  | 3.44% | 1.10% | 572 | 530 | 21044 | 24631 |
|  |  | rPS Medium<sub>Rapa</sub> |  | 3.38% | 0.81% | 290 | 125 | 9391 | 5596 |
|  |  | Oral Rapamycin |  | 5.00% | 2.78% | 409 | 277 | 13393 | 16075 |
|  |  | anti-CD3 mAb |  | 26.36% | 21.54% | 2778 | 2889 | 8523 | 3993 |
| A/B LN | CD8+ CD4-T cells | Non-Treated |  | 14.34% | 7.53% | 28754 | 39992 | 150281 | 136587 |
|  |  | Vehicle PS |  | 15.76% | 7.57% | 32666 | 37287 | 157804 | 145645 |
|  |  | rPS Medium<sub>Rapa</sub> |  | 16.80% | 8.99% | 14140 | 22110 | 57822 | 68078 |
|  |  | Oral Rapamycin |  | 13.33% | 7.05% | 17908 | 17034 | 106605 | 87909 |
|  |  | anti-CD3 mAb |  | 8.62% | 5.59% | 4361 | 6294 | 32656 | 35410 |
|  | CD8+ CD4+ (Double Positive) T cells | Non-Treated |  | 0.72% | 0.14% | 1223.8 | 1375.8 | 150281 | 136587 |
|  |  | Vehicle PS |  | 0.67% | 0.07% | 1059.6 | 990.9 | 157804 | 145645 |
|  |  | rPS Medium<sub>Rapa</sub> |  | 0.34% | 0.07% | 203 | 263.6 | 57822 | 68078 |
|  |  | Oral Rapamycin |  | 0.57% | 0.06% | 609 | 526.3 | 106605 | 87909 |
|  |  | anti-CD3 mAb |  | 0.59% | 0.13% | 210.6 | 229.4 | 32656 | 35410 |
|  | CD8- CD4+ T cells | Non-Treated |  | 81.93% | 6.52% | 117196 | 94999 | 150281 | 136587 |
|  |  | Vehicle PS |  | 81.40% | 6.96% | 121835 | 106895 | 157804 | 145645 |

|  |  |  |  |  |  |  |  |  |
| --- | --- | --- | --- | --- | --- | --- | --- | --- |
|  |  | rPS Medium<sub>Rapa</sub> | 79.42% | 7.73% | 41981 | 44469 | 57822 | 68078 |
|  |  | Oral Rapamycin | 83.50% | 5.90% | 86015 | 70025 | 106605 | 87909 |
|  |  | anti-CD3 mAb | 79.24% | 3.61% | 25765 | 27243 | 32656 | 35410 |
|  | CD8- CD4-<br>(Double Negative) | Non-Treated | 3.00% | 2.25% | 3108 | 1286 | 150281 | 136587 |
|  |  | Vehicle PS | 2.16% | 1.16% | 2244 | 1867 | 157804 | 145645 |
|  |  | rPS Medium<sub>Rapa</sub> | 3.46% | 1.55% | 1498 | 1374 | 57822 | 68078 |
|  |  | Oral Rapamycin | 2.62% | 1.28% | 2072 | 1357 | 106605 | 87909 |
|  |  | anti-CD3 mAb | 11.56% | 6.55% | 2320 | 1896 | 32656 | 35410 |

**Supplemental Table 10.** CD8+ T cell subsets overview in blood and A/B LN.

|  |  | Cell Type | Treatment | Mean Frequency of Cell Type+ of CD8+ T cells | Standard Deviation of Frequency of Cell Type+ of CD8+ T cells | Mean Count of Cell Type+ of CD8+ T cells | Standard Deviation of Count of Cell Type+ of CD8+ T cells | Mean Count of CD8+ T cells | Standard Deviation of Count of CD8+ T cells |
| --- | --- | --- | --- | --- | --- | --- | --- | --- | --- |
| Blood | CD38+ CD8+ T cells | Non-Treated |  | 1.24% | 1.70% | 16 | 9.8 | 4239 | 6122 |
|  |  | Vehicle PS |  | 0.56% | 0.15% | 26.2 | 18.9 | 5801 | 6339 |
|  |  | rPS Medium<sub>Rapa</sub> |  | 0.17% | 0.08% | 7 | 6 | 3535 | 2232 |
|  |  | Oral Rapamycin |  | 0.30% | 0.18% | 11.6 | 12.4 | 4229 | 4818 |
|  |  | anti-CD3 mAb |  | 0.73% | 0.29% | 8.2 | 4.9 | 1180 | 492 |
|  | CD73+ CD8+ T cells | Non-Treated |  | 29.06% | 3.53% | 1240 | 1734 | 4239 | 6122 |
|  |  | Vehicle PS |  | 26.16% | 3.29% | 1573 | 1775 | 5801 | 6339 |
|  |  | rPS Medium<sub>Rapa</sub> |  | 38.75% | 4.14% | 1438 | 1063 | 3535 | 2232 |
|  |  | Oral Rapamycin |  | 27.20% | 1.44% | 1143 | 1320 | 4229 | 4818 |
|  |  | anti-CD3 mAb |  | 31.34% | 4.02% | 372 | 165 | 1180 | 492 |
|  | FR4+ CD8+ T cells | Non-Treated |  | 16.32% | 1.51% | 748 | 1135 | 4239 | 6122 |
|  |  | Vehicle PS |  | 17.02% | 1.97% | 1077 | 1298 | 5801 | 6339 |
|  |  | rPS Medium<sub>Rapa</sub> |  | 7.02% | 2.32% | 254 | 169 | 3535 | 2232 |
|  |  | Oral Rapamycin |  | 14.32% | 1.82% | 622 | 715 | 4229 | 4818 |
|  |  | anti-CD3 mAb |  | 25.62% | 3.17% | 303 | 138 | 1180 | 492 |
|  | ICOS+ CD8+ T cells | Non-Treated |  | 0.82% | 0.69% | 12.8 | 7.79 | 4239 | 6122 |
|  |  | Vehicle PS |  | 0.45% | 0.41% | 14 | 8.09 | 5801 | 6339 |
|  |  | rPS Medium<sub>Rapa</sub> |  | 0.30% | 0.19% | 8.5 | 3.7 | 3535 | 2232 |
|  |  | Oral Rapamycin |  | 0.34% | 0.24% | 9.2 | 6.18 | 4229 | 4818 |
|  |  | anti-CD3 mAb |  | 0.75% | 0.52% | 7.8 | 3.56 | 1180 | 492 |
|  | Lag3+ CD8+ T cells | Non-Treated |  | 0.89% | 0.72% | 12.6 | 5.94 | 4239 | 6122 |
|  |  | Vehicle PS |  | 0.34% | 0.34% | 9.6 | 3.44 | 5801 | 6339 |
|  |  | rPS Medium<sub>Rapa</sub> |  | 0.37% | 0.19% | 10 | 0.82 | 3535 | 2232 |
|  |  | Oral Rapamycin |  | 0.41% | 0.28% | 9 | 5.24 | 4229 | 4818 |
|  |  | anti-CD3 mAb |  | 1.03% | 0.83% | 10 | 4.12 | 1180 | 492 |
|  | PD1+ CD8+ T cells | Non-Treated |  | 2.28% | 3.06% | 33.2 | 22.7 | 4239 | 6122 |
|  |  | Vehicle PS |  | 1.41% | 0.69% | 58.2 | 44.2 | 5801 | 6339 |
|  |  | rPS Medium<sub>Rapa</sub> |  | 0.44% | 0.21% | 18.5 | 19.2 | 3535 | 2232 |
|  |  | Oral Rapamycin |  | 1.02% | 0.59% | 27 | 14.3 | 4229 | 4818 |
|  |  | anti-CD3 mAb |  | 1.20% | 0.53% | 15.2 | 12.3 | 1180 | 492 |
|  | TIGIT+ CD8+ T cells | Non-Treated |  | 2.02% | 1.61% | 31 | 24.4 | 4239 | 6122 |
|  |  | Vehicle PS |  | 0.84% | 0.99% | 20.6 | 4.8 | 5801 | 6339 |

|  |  |  |  |  |  |  |  |  |
| --- | --- | --- | --- | --- | --- | --- | --- | --- |
|  |  | rPS Medium<sub>Rapa</sub> | 0.74% | 0.27% | 21.8 | 5 | 3535 | 2232 |
|  |  | Oral Rapamycin | 0.74% | 0.41% | 18.2 | 8.3 | 4229 | 4818 |
|  |  | anti-CD3 mAb | 1.76% | 1.36% | 17 | 6.2 | 1180 | 492 |
|  | CD25+<br>FoxP3+<br>CD8+<br>Tregs | Non-Treated | 0.72% | 0.65% | 9.4 | 4.72 | 4239 | 6122 |
|  |  | Vehicle PS | 0.27% | 0.22% | 7.4 | 1.52 | 5801 | 6339 |
|  |  | rPS Medium<sub>Rapa</sub> | 0.29% | 0.12% | 8.75 | 2.99 | 3535 | 2232 |
|  |  | Oral Rapamycin | 0.25% | 0.17% | 5.6 | 3.65 | 4229 | 4818 |
|  |  | anti-CD3 mAb | 0.68% | 0.55% | 6.6 | 2.61 | 1180 | 492 |
|  | CD38+<br>CD8+ T<br>cells | Non-Treated | 0.68% | 0.12% | 158 | 187.7 | 28754 | 39992 |
|  |  | Vehicle PS | 0.88% | 0.45% | 190.8 | 207.9 | 32666 | 37287 |
|  |  | rPS Medium<sub>Rapa</sub> | 0.23% | 0.18% | 25.8 | 45 | 14140 | 22110 |
|  |  | Oral Rapamycin | 0.54% | 0.26% | 68.3 | 62.7 | 17908 | 17034 |
|  |  | anti-CD3 mAb | 2.47% | 2.82% | 45.2 | 63.9 | 4361 | 6294 |
| A/B LN | CD73+<br>CD8+ T<br>cells | Non-Treated | 17.40% | 3.03% | 5883 | 8959 | 28754 | 39992 |
|  |  | Vehicle PS | 19.00% | 5.32% | 6498 | 7795 | 32666 | 37287 |
|  |  | rPS Medium<sub>Rapa</sub> | 27.20% | 5.77% | 4498 | 7444 | 14140 | 22110 |
|  |  | Oral Rapamycin | 18.80% | 6.37% | 3487 | 3247 | 17908 | 17034 |
|  |  | anti-CD3 mAb | 26.70% | 5.32% | 1045 | 1555 | 4361 | 6294 |
|  | FR4+<br>CD8+ T<br>cells | Non-Treated | 22.80% | 2.04% | 6380 | 8775 | 28754 | 39992 |
|  |  | Vehicle PS | 20.10% | 2.79% | 6118 | 6775 | 32666 | 37287 |
|  |  | rPS Medium<sub>Rapa</sub> | 10.60% | 5.80% | 1312 | 2199 | 14140 | 22110 |
|  |  | Oral Rapamycin | 19.00% | 3.87% | 2961 | 2673 | 17908 | 17034 |
|  |  | anti-CD3 mAb | 28.80% | 5.26% | 1223 | 1784 | 4361 | 6294 |
|  | ICOS+<br>CD8+ T<br>cells | Non-Treated | 1.60% | 0.73% | 256 | 178.57 | 28754 | 39992 |
|  |  | Vehicle PS | 1.55% | 1.25% | 188.4 | 178.87 | 32666 | 37287 |
|  |  | rPS Medium<sub>Rapa</sub> | 1.10% | 1.45% | 36.2 | 23.82 | 14140 | 22110 |
|  |  | Oral Rapamycin | 1.67% | 1.52% | 118 | 84.32 | 17908 | 17034 |
|  |  | anti-CD3 mAb | 3.61% | 2.47% | 61 | 65.37 | 4361 | 6294 |
|  | Lag3+<br>CD8+ T<br>cells | Non-Treated | 0.02% | 0.01% | 3.75 | 3.77 | 28754 | 39992 |
|  |  | Vehicle PS | 0.13% | 0.23% | 5.4 | 4.93 | 32666 | 37287 |
|  |  | rPS Medium<sub>Rapa</sub> | 0.56% | 1.15% | 6 | 4.18 | 14140 | 22110 |
|  |  | Oral Rapamycin | 0.13% | 0.21% | 4 | 1.83 | 17908 | 17034 |
|  |  | anti-CD3 mAb | 0.58% | 0.74% | 3.6 | 2.19 | 4361 | 6294 |
|  | PD1+<br>CD8+ T<br>cells | Non-Treated | 2.07% | 1.13% | 322.5 | 229.3 | 28754 | 39992 |
|  |  | Vehicle PS | 2.00% | 1.46% | 267 | 225.9 | 32666 | 37287 |
|  |  | rPS Medium<sub>Rapa</sub> | 2.19% | 3.06% | 70.2 | 53.2 | 14140 | 22110 |
|  |  | Oral Rapamycin | 1.65% | 0.98% | 176 | 144.7 | 17908 | 17034 |
|  |  | anti-CD3 mAb | 7.60% | 6.55% | 106.2 | 94.6 | 4361 | 6294 |
|  | TIGIT+<br>CD8+ T<br>cells | Non-Treated | 0.25% | 0.22% | 31.5 | 23.2 | 28754 | 39992 |
|  |  | Vehicle PS | 0.41% | 0.44% | 22.8 | 17.5 | 32666 | 37287 |
|  |  | rPS Medium<sub>Rapa</sub> | 0.46% | 0.77% | 10.8 | 6.9 | 14140 | 22110 |
|  |  | Oral Rapamycin | 0.57% | 0.59% | 28 | 22.4 | 17908 | 17034 |
|  |  | anti-CD3 mAb | 2.37% | 2.93% | 17.8 | 11.8 | 4361 | 6294 |
|  | CD25+<br>FoxP3+<br>CD8+<br>Tregs | Non-Treated | 0.05% | 0.03% | 9.75 | 10.24 | 28754 | 39992 |
|  |  | Vehicle PS | 0.16% | 0.22% | 11.4 | 11.08 | 32666 | 37287 |
|  |  | rPS Medium<sub>Rapa</sub> | 0.21% | 0.32% | 6 | 4.06 | 14140 | 22110 |
|  |  | Oral Rapamycin | 0.22% | 0.35% | 8.75 | 6.24 | 17908 | 17034 |
|  |  | anti-CD3 mAb | 1.00% | 1.52% | 6.6 | 5.41 | 4361 | 6294 |

**Supplementary Table 11.** CD4+ T cell subsets overview in blood.

| Cell Type | Treatment | Mean Frequency of Cell Type+ of CD4+ T cells | Standard Deviation of Frequency of Cell Type+ of CD4+ T cells | Mean Count of Cell Type+ of CD4+ T cells | Standard Deviation of Count of Cell Type+ of CD4+ T cells | Mean Count of CD4+ T cells | Standard Deviation of Count of CD4+ T cells |
| --- | --- | --- | --- | --- | --- | --- | --- |
| CD38+ CD4+ T cells | Non-Treated | 0.15% | 0.08% | 11.4 | 14.8 | 11080 | 16769 |
|  | Vehicle PS | 0.26% | 0.10% | 38 | 50 | 14640 | 17729 |
|  | rPS Medium<sub>Rapa</sub> | 0.16% | 0.07% | 8 | 5.4 | 5556 | 3362 |
|  | Oral Rapamycin | 0.21% | 0.12% | 13 | 13.5 | 8740 | 10970 |
|  | anti-CD3 mAb | 0.32% | 0.23% | 16.4 | 14.7 | 4554 | 1844 |
| CD73+ CD4+ T cells | Non-Treated | 14.10% | 2.65% | 1294 | 1703 | 11080 | 16769 |
|  | Vehicle PS | 11.80% | 3.04% | 1861 | 2550 | 14640 | 17729 |
|  | rPS Medium<sub>Rapa</sub> | 15.59% | 4.44% | 949 | 719 | 5556 | 3362 |
|  | Oral Rapamycin | 11.60% | 4.60% | 790 | 852 | 8740 | 10970 |
|  | anti-CD3 mAb | 20.14% | 4.04% | 940 | 507 | 4554 | 1844 |
| FR4+ CD4+ T cells | Non-Treated | 42.10% | 4.42% | 4633 | 7035 | 11080 | 16769 |
|  | Vehicle PS | 41.48% | 2.60% | 6076 | 7410 | 14640 | 17729 |
|  | rPS Medium<sub>Rapa</sub> | 27.85% | 5.99% | 1644 | 1177 | 5556 | 3362 |
|  | Oral Rapamycin | 38.40% | 3.91% | 3142 | 3782 | 8740 | 10970 |
|  | anti-CD3 mAb | 59.50% | 2.94% | 2709 | 1138 | 4554 | 1844 |
| ICOS+ CD4+ T cells | Non-Treated | 3.49% | 4.08% | 143.8 | 147.54 | 11080 | 16769 |
|  | Vehicle PS | 2.39% | 1.01% | 276.8 | 270.98 | 14640 | 17729 |
|  | rPS Medium<sub>Rapa</sub> | 0.98% | 0.45% | 63.75 | 59.06 | 5556 | 3362 |
|  | Oral Rapamycin | 3.85% | 3.68% | 179.6 | 132.32 | 8740 | 10970 |
|  | anti-CD3 mAb | 4.41% | 2.01% | 205.4 | 158.48 | 4554 | 1844 |
| Lag3+ CD4+ T cells | Non-Treated | 0.01% | 0.01% | 0.6 | 0.89 | 11080 | 16769 |
|  | Vehicle PS | 0.00% | 0.00% | 0.2 | 0.45 | 14640 | 17729 |
|  | rPS Medium<sub>Rapa</sub> | 0.02% | 0.03% | 0.75 | 0.96 | 5556 | 3362 |
|  | Oral Rapamycin | 0.00% | 0.00% | 0.6 | 1.34 | 8740 | 10970 |
|  | anti-CD3 mAb | 0.00% | 0.01% | 0.2 | 0.45 | 4554 | 1844 |
| PD1+ CD4+ T cells | Non-Treated | 2.97% | 2.60% | 132.2 | 106.5 | 11080 | 16769 |
|  | Vehicle PS | 1.99% | 0.73% | 261.6 | 300 | 14640 | 17729 |
|  | rPS Medium<sub>Rapa</sub> | 1.35% | 0.40% | 83.5 | 68.3 | 5556 | 3362 |
|  | Oral Rapamycin | 2.80% | 2.28% | 130.4 | 96.5 | 8740 | 10970 |
|  | anti-CD3 mAb | 4.28% | 0.98% | 197.4 | 108.4 | 4554 | 1844 |
| TIGIT+ CD4+ T cells | Non-Treated | 0.18% | 0.19% | 9.6 | 9.8 | 11080 | 16769 |
|  | Vehicle PS | 0.08% | 0.04% | 7.8 | 6.4 | 14640 | 17729 |
|  | rPS Medium<sub>Rapa</sub> | 0.11% | 0.05% | 6.3 | 4.3 | 5556 | 3362 |
|  | Oral Rapamycin | 0.16% | 0.16% | 7.2 | 5.2 | 8740 | 10970 |
|  | anti-CD3 mAb | 0.08% | 0.04% | 3 | 1.6 | 4554 | 1844 |
| CD25+<br>FoxP3+<br>CD4+<br>Tregs | Non-Treated | 0.30% | 0.25% | 25 | 37 | 11080 | 16769 |
|  | Vehicle PS | 0.23% | 0.16% | 49.6 | 80.78 | 14640 | 17729 |
|  | rPS Medium<sub>Rapa</sub> | 0.22% | 0.25% | 15 | 21.49 | 5556 | 3362 |
|  | Oral Rapamycin | 0.24% | 0.09% | 16.4 | 15.73 | 8740 | 10970 |
|  | anti-CD3 mAb | 0.46% | 0.40% | 17 | 8.15 | 4554 | 1844 |
| CD44hi/lo<br>FoxP3+<br>CD4+<br>Tregs | Non-Treated | 3.33% | 4.18% | 257 | 346.66 | 11080 | 16769 |
|  | Vehicle PS | 1.73% | 0.78% | 440.6 | 562.05 | 14640 | 17729 |
|  | rPS Medium<sub>Rapa</sub> | 1.09% | 0.58% | 172.5 | 150.58 | 5556 | 3362 |
|  | Oral Rapamycin | 3.29% | 3.63% | 213.8 | 149.83 | 8740 | 10970 |
|  | anti-CD3 mAb | 3.45% | 1.49% | 273.2 | 176.42 | 4554 | 1844 |
| CD44-<br>FoxP3-<br>CD4+ T<br>cells | Non-Treated | 91.20% | 6.56% | 10522.8 | 16056.81 | 11080 | 16769 |
|  | Vehicle PS | 93.82% | 1.88% | 13715.8 | 16518.45 | 14640 | 17729 |
|  | rPS Medium<sub>Rapa</sub> | 94.65% | 2.54% | 5247.75 | 3179.22 | 5556 | 3362 |
|  | Oral Rapamycin | 91.90% | 5.27% | 8313 | 10629.21 | 8740 | 10970 |
|  | anti-CD3 mAb | 87.72% | 4.89% | 4005.4 | 1600.28 | 4554 | 1844 |
| CD44+<br>FoxP3-<br>CD4+<br>TEM | Non-Treated | 3.72% | 1.59% | 271.4 | 357.65 | 11080 | 16769 |
|  | Vehicle PS | 3.06% | 1.02% | 443.4 | 599.53 | 14640 | 17729 |
|  | rPS Medium<sub>Rapa</sub> | 2.05% | 0.49% | 119.5 | 81.72 | 5556 | 3362 |
|  | Oral Rapamycin | 3.48% | 1.70% | 205 | 199.32 | 8740 | 10970 |
|  | anti-CD3 mAb | 5.92% | 2.40% | 268 | 176.41 | 4554 | 1844 |
| CD44+<br>FoxP3-<br>Tanergic | Non-Treated | 0.34% | 0.30% | 23.2 | 19.55 | 11080 | 16769 |
|  | Vehicle PS | 0.21% | 0.15% | 36.6 | 60.77 | 14640 | 17729 |
|  | rPS Medium<sub>Rapa</sub> | 0.26% | 0.19% | 13.75 | 15.65 | 5556 | 3362 |
|  | Oral Rapamycin | 0.12% | 0.12% | 6.4 | 5.94 | 8740 | 10970 |
|  | anti-CD3 mAb | 0.19% | 0.15% | 7 | 2.83 | 4554 | 1844 |
| CD38+ CD4+ T cells | Non-Treated | 1.36% | 0.44% | 1365.5 | 229.3 | 117196 | 94999 |
|  | Vehicle PS | 1.31% | 0.35% | 1391.8 | 225.9 | 121835 | 106895 |
|  | rPS Medium<sub>Rapa</sub> | 0.69% | 0.30% | 260 | 53.2 | 41981 | 44469 |
|  | Oral Rapamycin | 1.11% | 0.34% | 826.8 | 144.7 | 86015 | 70025 |
|  | anti-CD3 mAb | 2.89% | 0.68% | 622.2 | 94.6 | 25765 | 27243 |
| CD73+ CD4+ T cells | Non-Treated | 13.10% | 1.22% | 14655 | 10417 | 117196 | 94999 |
|  | Vehicle PS | 13.50% | 1.52% | 15644 | 13687 | 121835 | 106895 |
|  | rPS Medium<sub>Rapa</sub> | 18.96% | 3.52% | 7458 | 7609 | 41981 | 44469 |
|  | Oral Rapamycin | 15.23% | 3.57% | 11347 | 7702 | 86015 | 70025 |
|  | anti-CD3 mAb | 29.50% | 7.90% | 6310 | 6151 | 25765 | 27243 |
| FR4+ CD4+ T cells | Non-Treated | 44.05% | 5.77% | 53641 | 48671 | 117196 | 94999 |
|  | Vehicle PS | 41.84% | 2.55% | 50494 | 43611 | 121835 | 106895 |
|  | rPS Medium<sub>Rapa</sub> | 32.76% | 5.18% | 13518 | 14606 | 41981 | 44469 |
|  | Oral Rapamycin | 41.13% | 2.71% | 34476 | 27085 | 86015 | 70025 |
|  | anti-CD3 mAb | 59.98% | 5.28% | 16013 | 17933 | 25765 | 27243 |
| ICOS+ CD4+ T cells | Non-Treated | 5.78% | 2.38% | 5352.75 | 1667.16 | 117196 | 94999 |
|  | Vehicle PS | 5.91% | 3.20% | 4621.6 | 3398.2 | 121835 | 106895 |
|  | rPS Medium<sub>Rapa</sub> | 6.16% | 3.85% | 1658.4 | 939.31 | 41981 | 44469 |
|  | Oral Rapamycin | 6.24% | 3.83% | 3591.5 | 2230.53 | 86015 | 70025 |
|  | anti-CD3 mAb | 18.36% | 11.45% | 3035 | 2522 | 25765 | 27243 |
| Lag3+ CD4+ T cells | Non-Treated | 0.14% | 0.18% | 111.25 | 101.16 | 117196 | 94999 |
|  | Vehicle PS | 1.83% | 4.00% | 60.6 | 34.93 | 121835 | 106895 |
|  | rPS Medium<sub>Rapa</sub> | 0.15% | 0.25% | 54.2 | 74.65 | 41981 | 44469 |
|  | Oral Rapamycin | 0.03% | 0.01% | 26.75 | 23.68 | 86015 | 70025 |
|  | anti-CD3 mAb | 0.08% | 0.08% | 26.6 | 36.2 | 25765 | 27243 |
| PD1+ CD4+ T cells | Non-Treated | 5.52% | 2.02% | 5367.8 | 2337.4 | 117196 | 94999 |
|  | Vehicle PS | 5.92% | 2.44% | 5313.6 | 4188.6 | 121835 | 106895 |
|  | rPS Medium<sub>Rapa</sub> | 6.87% | 3.96% | 1967.8 | 1279.5 | 41981 | 44469 |
|  | Oral Rapamycin | 6.37% | 2.72% | 4196.3 | 2765.6 | 86015 | 70025 |
|  | anti-CD3 mAb | 15.52% | 5.24% | 3044.2 | 2635.8 | 25765 | 27243 |
| TIGIT+ CD4+ T cells | Non-Treated | 0.53% | 0.35% | 779 | 293.7 | 117196 | 94999 |
|  | Vehicle PS | 1.78% | 2.47% | 741.6 | 583.9 | 121835 | 106895 |
|  | rPS Medium<sub>Rapa</sub> | 1.28% | 0.55% | 401.4 | 229.7 | 41981 | 44469 |
|  | Oral Rapamycin | 0.87% | 0.41% | 610 | 474.9 | 86015 | 70025 |
|  | anti-CD3 mAb | 1.69% | 0.72% | 372.6 | 322.8 | 25765 | 27243 |
| CD25+ FoxP3+ CD4+ Tregs | Non-Treated | 2.26% | 1.01% | 2636.75 | 2406.36 | 117196 | 94999 |
|  | Vehicle PS | 2.65% | 1.13% | 2503.6 | 2422.64 | 121835 | 106895 |
|  | rPS Medium<sub>Rapa</sub> | 2.88% | 0.51% | 1194 | 1249.92 | 41981 | 44469 |
|  | Oral Rapamycin | 2.75% | 0.51% | 2428 | 1936.12 | 86015 | 70025 |
|  | anti-CD3 mAb | 4.37% | 1.32% | 1080.2 | 1045.17 | 25765 | 27243 |
| CD44hi/lo FoxP3+ CD4+ Tregs | Non-Treated | 6.49% | 2.73% | 9140.5 | 6105.17 | 117196 | 94999 |
|  | Vehicle PS | 7.05% | 3.53% | 8426.2 | 7065.51 | 121835 | 106895 |
|  | rPS Medium<sub>Rapa</sub> | 7.74% | 3.14% | 3495.6 | 2709.95 | 41981 | 44469 |
|  | Oral Rapamycin | 7.79% | 2.95% | 7747.25 | 5453.71 | 86015 | 70025 |
|  | anti-CD3 mAb | 17.64% | 7.77% | 4710.2 | 4112.23 | 25765 | 27243 |
| CD44- FoxP3- CD4+ T cells | Non-Treated | 85.33% | 4.09% | 101418 | 84755.01 | 117196 | 94999 |
|  | Vehicle PS | 84.52% | 6.30% | 107920.2 | 96921.73 | 121835 | 106895 |
|  | rPS Medium<sub>Rapa</sub> | 85.80% | 5.35% | 37274.8 | 40903.05 | 41981 | 44469 |
|  | Oral Rapamycin | 83.55% | 5.57% | 74550 | 62403.51 | 86015 | 70025 |
|  | anti-CD3 mAb | 67.32% | 11.77% | 19292.6 | 21683.95 | 25765 | 27243 |
| CD44+ FoxP3- CD4+ TEM | Non-Treated | 5.04% | 1.00% | 5613.25 | 4111.87 | 117196 | 94999 |
|  | Vehicle PS | 3.99% | 0.82% | 4256.8 | 3373.02 | 121835 | 106895 |
|  | rPS Medium<sub>Rapa</sub> | 2.59% | 1.49% | 816.6 | 711.84 | 41981 | 44469 |
|  | Oral Rapamycin | 4.62% | 2.12% | 2952.5 | 1801.94 | 86015 | 70025 |
|  | anti-CD3 mAb | 7.98% | 4.44% | 1490.4 | 1378.81 | 25765 | 27243 |
| CD44+ FoxP3- Tanergic | Non-Treated | 0.89% | 0.41% | 921.5 | 520.88 | 117196 | 94999 |
|  | Vehicle PS | 0.99% | 0.18% | 1145.6 | 952.84 | 121835 | 106895 |
|  | rPS Medium<sub>Rapa</sub> | 1.16% | 0.58% | 364.4 | 271.49 | 41981 | 44469 |
|  | Oral Rapamycin | 0.97% | 0.33% | 707.5 | 559.21 | 86015 | 70025 |
|  | anti-CD3 mAb | 1.04% | 0.19% | 250 | 259.65 | 25765 | 27243 |
